# GFPrint™: A MACHINE LEARNING TOOL FOR TRANSFORMING GENETIC DATA INTO CLINICAL INSIGHTS

**DOI:** 10.1101/2024.03.08.584090

**Authors:** Guillermo Sanz-Martín, Daniela Paula Migliore, Pablo Gómez del Campo, José del Castillo-Izquierdo, Juan Manuel Domínguez

## Abstract

The increasing availability of massive genetic sequencing data in the clinical setting has triggered the need for appropriate tools to help fully exploit the wealth of information these data possess. GFPrint^™^ is a proprietary streaming algorithm designed to meet that need. By extracting the most relevant functional features, GFPrint^™^ transforms high-dimensional, noisy genetic sequencing data into an embedded representation, allowing unsupervised models to create data clusters that can be re-mapped to the original clinical information. Ultimately, this allows the identification of genes and pathways relevant to disease onset and progression. GFPrint^™^ has been tested and validated using two cancer genomic datasets publicly available. Analysis of the TCGA dataset has identified panels of genes whose mutations appear to negatively influence survival in non-metastatic colorectal cancer (15 genes), epidermoid non-small cell lung cancer (167 genes) and pheochromocytoma (313 genes) patients. Likewise, analysis of the Broad Institute dataset has identified 75 genes involved in pathways related to extracellular matrix reorganization whose mutations appear to dictate a worse prognosis for breast cancer patients. GFPrint^™^ is accessible through a secure web portal and can be used in any therapeutic area where the genetic profile of patients influences disease evolution.

## INTRODUCTION

Over the last decades, advances in cancer therapy have contributed to the 33% reduction in mortality rate observed between 1991 and 2020 [1]. The greatest contribution to these better outcomes came from the molecular characterization of tumors [2-4], influenced by global initiatives like the Cancer Genome Project [5] and the complete whole genome sequencing of tumors carried out by the Pan-Cancer Analysis of Whole Genomes (PCAWG) [6, 7]. This has led to an increased understanding of the molecular pathways that underlie cancer, favoring the discovery of molecular prognostic biomarkers and the incorporation of novel targeted agents. Nonetheless, the incidence of cancer in the global population is still high [8, 9] and is expected to remain high in coming years for several reasons, the most relevant being population aging associated to an increasing life expectancy [10]. Moreover, for most patients with metastatic cancer, the duration of clinical benefit is limited and is followed by drug resistance and cancer progression, imposing novel challenges in cancer treatment [11, 12]. Consequently, to continue extending survival and improving the patients’ quality of life, research endeavors to better understand the molecular physiopathology of cancer must continue.

In the last few years, genomic sequencing technologies have experienced a staggering evolution that has led to a cost reduction of nearly 7 orders of magnitude concomitant with a speed increase of 4 orders of magnitude compared to the first technologies that were used for the Human Genome Project [13, 14]. Therefore, obtaining comprehensive genetic information from cancer patients is expected to become routine in the near term. Ideally, this will transform the current landscape of precision oncology and will likely cause a boost in tissue-agnostic treatments dictated by the genetic profile of the patient rather than by the tissue where the disease originated. This will make imperative the use of appropriate tools to handle, interpret and exploit the massive amount of information that will become available to maximize its impact. Currently available tools, such as genome-wide complex trait analysis (GCTA) [15], polygenic risk score (PRS) [16] and the like, appear insufficient to that end.

Artificial intelligence (AI) and machine learning (ML) tools are expected to fill this gap given their unprecedented ability to integrate multiple types of data, extracting relevant features that can inform on hidden causalities, enable accurate predictions, build useful classifications, and generate novel information [17]. Different ML tools have been developed to leverage genetic information of patients for different purposes within a precision medicine context (e.g., patients’ stratification, disease risk assessment or identification of susceptibility features) [18, 19]. All of them utilize classic ML algorithms and are intended for specific applications. We have developed a novel ML tool named Genetic Fingerprint (GFPrint^™^), which offers added benefits over existing tools, as will be discussed later. GFPrint^™^ is designed to exploit genomic information from patients for prognostic and therapeutic purposes. GFPrint^™^ analyzes sequencing data to create a synthetic representation of patients in a latent space; this is combined with clinical information from the same patients to obtain conclusions that can be used to propose unexplored biomarkers, predict possible outcomes, identify novel therapeutic targets, or identify patterns that may help selecting the most appropriate treatment for each patient. To validate GFPrint^™^, a retrospective analysis has been run with data from cancer patients available in datasets whose content is public and whose size was large enough to enable a broad study pursuing two main goals: to evaluate the performance of GFPrint^™^ and to assess its ability to deliver useful insights that can be leveraged in novel therapeutic approaches. This paper summarizes the results obtained.

## METHODS

### Study datasets

Two different datasets were used in this study. The first dataset corresponded to DNA whole exome sequence (WES) data in The Cancer Genome Atlas (TCGA) pan-cancer database [20, 21]. The data were originally obtained from tumor samples at diagnosis from 11,811 patients and mutations were annotated with respect to the human genome assembly HG38. More information about sample types and DNA sequencing methodology can be found at https://gdc.cancer.gov/. The second dataset corresponded to DNA WES from the Broad Institute of MIT and Harvard. The data were originally obtained from tumor samples at diagnosis from 945 patients and mutations were annotated with respect to the human genome assembly HG19. To make it consistent with the previous dataset, they were processed with the UCSC liftOver software so that they were finally annotated with respect to the HG38 assembly. From all the associated clinical information available in both datasets, only overall survival (OS) was considered.

### Statistical analysis

All statistical analyses were performed using R version 4.3.2 and the corresponding packages therein as described below. Comparisons between categorical variables were based on the χ2 test while comparisons between continuous variables were based on the Mann-Whitney U test. Tests were performed with stats base package. According to TCGA definition, OS was calculated from the date of diagnosis to the date of death from any cause or to the last follow-up examination (censored event). OS curves were calculated according to the Kaplan-Meier method and compared using the log-rank test before and after stratification according to clinical variables expected to play an important prognostic role in cancer. Multivariate analysis was performed using the Cox proportional-hazard model using time-dependent co-variables that enabled quantification of the existence of interactions between biological variables and time. Survival analysis was performed using survival package version 3.5-7. Unless otherwise explained, differences were considered significant if the p-value was < .05 (two sided).

### GFPrint^™^

GFPrint^™^ is a DNA sequence encoder based on a proprietary streaming algorithm. As input, it receives a .csv file in which each row represents a base mutation, and columns contains the anonymized patient ID, the chromosome number, the position of the mutation, the reference allele, and the mutated allele. The expected runtime for each experiment depends on the total amount of mutations, considering an approximate rate of 1E4 mutations analyzed per minute (e.g., 100 minutes runtime for analyzing 1 million mutations). GFPrint^™^ can be accessed upon request at. The encoding of dataset from TCGA (11,811 patients with a combined total amount of 3 million mutations) took about 3 hours and 45 minutes to be totally processed. The system outputs a .csv file containing for each patient a DNA fingerprint compiling the uploaded information in a vector of 10,240 positions with a 2% sparsity. These fingerprints are used for further machine learning and statistical analysis. To extract nonlinear relationships of the data, we used unsupervised learning and dimensionality reduction techniques such as UMAP, from the umap-learn package version 0.5.3, to place the genetic embeddings in a three-dimensional space. The space was then clustered using the DBSCAN [22] algorithm from the scikit-learn package version 1.2.2. The Python version used was 3.10.9.

### Pathway enrichment analysis

Pathway enrichment analysis (PEA) was conducted using clusterProfiler 4.3.8 [23] package in R version 4.3.2, a powerful tool designed for statistical analysis and visualization of functional profiles for genes and gene clusters. Mutations were annotated to their corresponding genes with Variant effect predictor (VEP) [24] software. We utilized the enrichKEGG function to perform KEGG pathway enrichment analysis, enrichGO for Gene Otnhology, enrichWP for Wikipathways and for the analysis of Reactome we used the function enrichPathway from the package ReactomePA [25]. In all cases, the adjusted p-value was calculated as false discovery rate (FDR) by using the Benjamini-Hochberg procedure [26].

## RESULTS

### Generation of a latent space with data from The Cancer Genome Atlas

To test the performance of GFPrint^™^, clinical and whole exome sequencing (WES) data from The Cancer Genome Atlas (TCGA) database were analyzed. This dataset is the largest repository of genomic data from cancer patients whose content is publicly available, and this is the reason why it was selected. The data for this analysis corresponded to tumors of 145 histotypes obtained from 11,811 patients. Sequences were encrypted using the GFPrint^™^ encoding system and the embedded information was processed by the GFPrint^™^ framework to create a latent space (Fig. 1A) in which each point corresponds to the WES of one individual tumor and can therefore be considered as a virtual representation of a patient’s tumor. After generating the latent space, the tool was able to identify 3 different clusters of patients, different from each other according to their genetic features extracted by GFPrint^™^.

**Figure 1:**
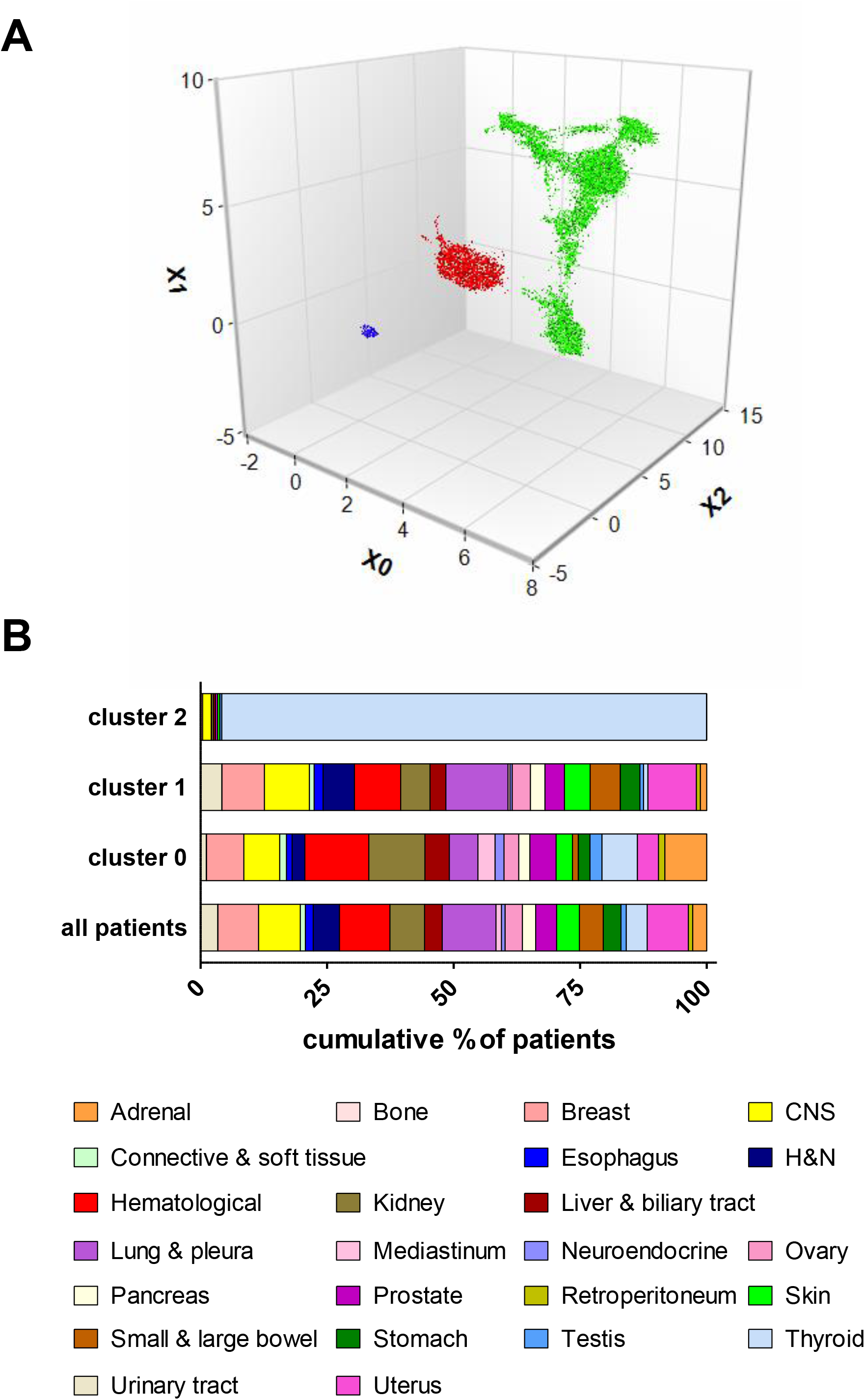
Analysis of the TCGA WES dataset with GFPrint^™^. **A**: Latent space for the TCGA WES dataset. The dataset, available through the Cancer Genome Consortium, was processed as described in the text. The three major clusters identified are denoted in different colors: cluster 0 (N = 2,681) in red, cluster 1 (N = 8,935) in green and cluster 2 (N= 236) in blue. **B**:Distribution of cancer groups by cluster. 25 cancer groups were created as described in the text, and their distribution in each cluster according to their percentage of patients is shown. CNS, central nervous system; H&N, head and neck.

To explore the resulting clusters and gain insights on their clinical meaning, differences in the outcome of these patients, expressed as overall survival (OS), were analyzed. The dataset was split in 25 different cancer groups according to the primary diagnosis as specified in the TCGA dataset (Supplementary Table 1). One of these 25 cancer groups (spermatic cord) included only one patient, so it was excluded from the analysis. Figure 1B shows the distribution of the remaining 24 cancer groups within each cluster. Notably, whilst clusters 0 and 1 showed an unbiased distribution of patients within the different cancer groups, cluster 2 was almost entirely populated with thyroid cancer patients (226 cases out of 236, i.e., 96%). Therefore, any conclusion regarding the clinical outcome of this cluster will be likely driven by the cancer type rather than by genetic features, and consequently this cluster was discarded for subsequent analysis.

**Table 1:** Pathologies in which cluster assignment had a significant influence in the clinical outcome according to the multivariate analysis. In this case, a p < .1 threshold in MOS differences between both clusters was set to determine the statistical significance. n, number of patients in the cluster; HR, hazard ratio; CI95, 95% confidence interval; CRC, colorectal cancer; eNSCLC, epidermoid non-small cell lung cancer.

| Pathology | p-value | HR | HR CI95 | n |
| --- | --- | --- | --- | --- |
| CRC | .027 | 2.294 | 1.1-4.786 | 545 |
| Melanoma | .009 | 2.156 | 1.215-3.827 | 452 |
| Mouth cancer | .064 | 2.179 | 0.957-4.961 | 174 |
| eNSCLC | .068 | 1.509 | 0.97-2.347 | 574 |
| Ovarian cancer | .049 | 1.366 | 1.001-1.864 | 99 |
| Pheochromocytoma | .037 | 9.102 | 1.141-72.590 | 147 |
| Pleura | .089 | 1.583 | 0.932-2.687 | 76 |
| Urothelial carcinoma | .047 | 1.677 | 1.007-2.792 | 397 |

When analyzing the differences in clinical outcome between the remaining two clusters for the 24 cancer groups, significant OS differences (p < .05 in median overall survival, MOS) were observed for 6 of the 24 groups (Fig. 2 and Suppl. Table 2). Three of these (adrenal, CNS and mediastinum) showed longer survival for patients in cluster 0 while the other three (small and large bowel, lung and pleura, and urinary tract) showed longer survival for patients in cluster 1, suggesting that cluster assignment (hence genetic features) may be linked to differences in the clinical outcome. After this first insight, all the 24 cancer groups were investigated further in the search for relevant patterns that were not easily observable at a glance because of the heterogeneity of the groups in terms of histopathology. Therefore, cancer groups were subdivided into different pathologies which were studied separately (the individual pathologies for each of the 24 groups are described in Suppl. Table 3). Then, for each pathology, 4 confounders additional to cluster assignment were considered: sex, age, disease stage and the presence or absence of metastasis. These 5 variables were used to run stratification analysis as well as statistical multivariate analysis to weight out the individual effect of each variable on the clinical outcome after adjusting for the effects of the other variables. There were 8 pathologies in which cluster assignment had a statistically significant influence in the clinical outcome (Table 1). The stratification analysis showed that only 5 of these 8 pathologies showed interaction between cluster assignment and one of the additional confounders, particularly disease stage or metastatic status. These 5 cases were colorectal cancer (n=545), epidermoid non-small cell lung cancer (n=574), pheochromocytoma (n=147), urothelial carcinoma (n=397) and melanoma (n=452).

**Figure 2:**
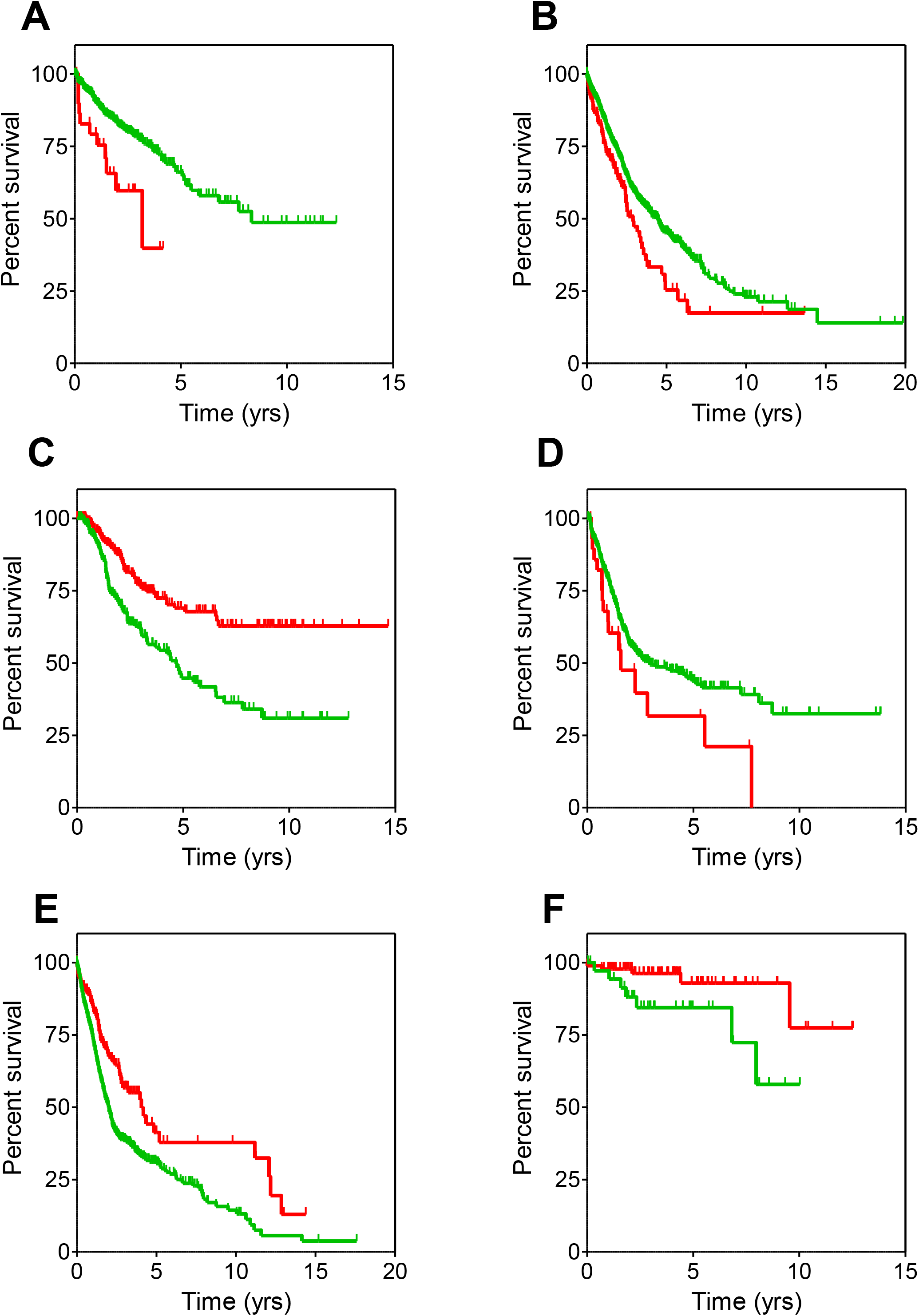
Survival curves for cancer groups. Survival curves for the 6 cancer groups where statistically significant differences (p < .05) between cluster 0 (red) and cluster 1 (green) patients were observed. Resulting values for MOS and hazard ratio are shown in Suppl. Table 2. **A**: small and large bowel; **B**: lung and pleura; **C**: adrenal; **D**: urinary tract; **E**: CNS; **F**: mediastinum. Censored patients are denoted with spikes. Kaplan-Meier curves were built using the clinical data available in the TCGA dataset in R.

### Colorectal cancer

As observed in Figure 2A, longer survival was observed for patients with large and small bowel cancers belonging to cluster 1 compared to those belonging to cluster 0. This result was related to differential OS in the colorectal cancer (CRC) subgroup (Fig. 3A). Indeed, CRC patients in cluster 0 (n=28) showed a significantly shorter OS compared to CRC patients in cluster 1 (n=517) (MOS, 3.1 years (CI95, 1.49-not reached, NR) vs. 8.3 years (CI95, 5.84-NR) for cluster 0 and cluster 1, respectively; hazard ratio (HR), 2.5 (CI95, 1.28-4.76); p-value=.007). Likewise, in multivariate analysis, shorter OS was significantly correlated to cluster 0 (p-value=.027 for MOS, see Table 1).

**Figure 3:**
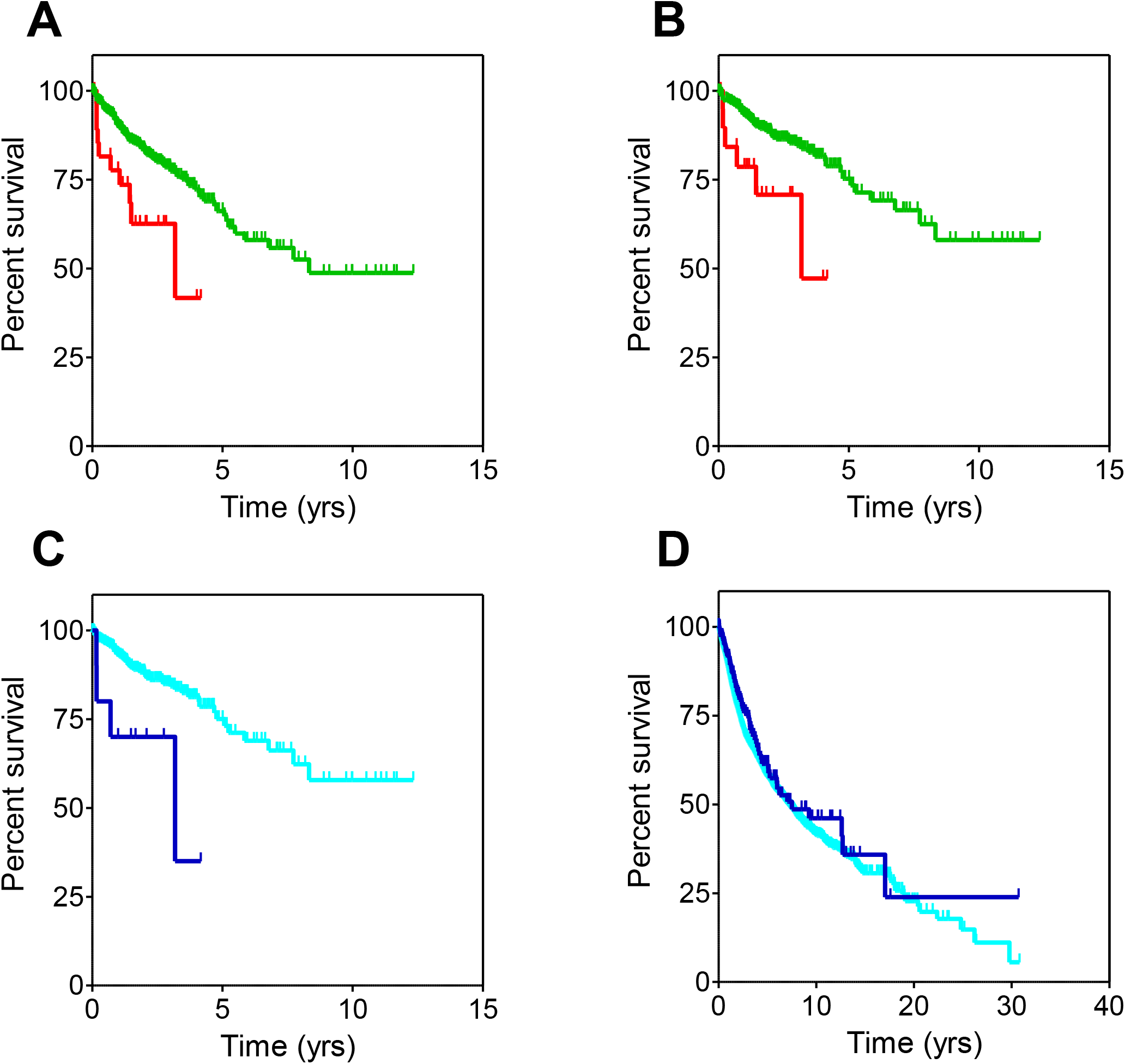
Overall survival curves for CRC patients. **A**: all CRC patients sorted by cluster. **B**: non-metastatic CRC patients sorted by cluster. **C**: non-metastatic CRC patients sorted by the presence or absence of mutations in any of the 15 genes described in the text. **D**: all patients included in the TCGA dataset sorted by the presence or absence of mutations in any of the above genes. Censored patients are denoted with spikes. Color codes are the same for all panels: red for cluster 0, green for cluster 1, dark blue for patients with mutations and light blue for patients without mutations in the intended genes.

Notably, the stratification analysis revealed that cluster assignment influenced survival only in CRC patients with non-metastatic disease. Indeed, Figure 3B shows that patients belonging to cluster 0 (n=20) presented a significantly shorter OS than those in cluster 1 (n=424) (MOS, 3.1 years (CI95, 3.1-NR) and NR (CI95, 8.3-NR) for cluster 0 and cluster 1 respectively; HR, 2.94 (CI95, 1.27-6.67); p-value=.009).

The results above suggested that the genetic profile of non-metastatic CRC patients in cluster 0 may be influencing their worse disease progression. The mutational landscape of these patients was then studied, and it turned out that the mutation burden of patients with poorer survival rate was smaller: patients belonging to cluster 0 showed mutations in 2,323 genes, whilst those belonging to cluster 1 showed mutations in 19,181 genes. Of these, 2,308 genes were mutated in both clusters, therefore a small set of only 15 genes (shown in Suppl. Table 4) were mutated exclusively in patients assigned to cluster 0. We then evaluated whether mutations in these genes could be used as surrogate markers for cluster 0, replacing the need for WES analysis to identify those patients with poor prognosis. As shown in Figure 3C, non-metastatic CRC patients with mutations in any of the 15 genes (n=10) showed significantly worse OS (MOS, 3.2 years; CI95, 0.7-NR) than those presenting no mutations (n=434) (MOS, NR; CI95, 8.3-NR; HR, 3.57; CI95, 1.28-10.0; p-value=.01). Interestingly, when mutations in these 15 genes were evaluated along the entire TCGA dataset regardless their cancer origin, no significant differences in OS were observed (HR, 0.83; CI95, 0.67-1.04; p-value=.1) (Fig. 3D).

In view of these results, it is tempting to hypothesize that mutations in these 15 genes may be influencing the worse evolution of non-metastatic CRC patients and therefore they may constitute a relevant panel of biomarkers to predict worse prognosis for this type of patients. Clearly, more experimental work is needed to obtain further evidence to support this point as well as to determine if they deserve to be investigated as possible novel targets of therapeutic intervention.

### Epidermoid non-small cell lung cancer

Figure 2B showed that patients belonging to cluster 0 within the “lung and pleura” cancer group presented a worse disease outcome, with a significantly shorter median OS than those included in cluster 1. However, when this wide group was subdivided into the different pathologies encompassed in it (adenocarcinoma non-small cell lung cancer (aNSCLC), epidermoid non-small cell lung cancer (eNSCLC) and mesothelioma), such differences remained only for stage I eNSCLC patients (n=266) (Figure 4A). In this clinical subset, patients belonging to cluster 0 (n=29) had significantly shorter OS compared to those of included in cluster 1 (n=237). MOS were 2.5 (CI95, 1.17-NR) and 5.4 years (CI95, 4.75-7.34) for cluster 0 and cluster 1, respectively (HR, 2.2; CI95, 1.32-3.85; p-value=.003). Moreover, the prognosis of stage I eNSCLC patients belonging to cluster 0 is comparable to patients diagnosed at stage IV (n=8; MOS, 2.0 years; CI95 1.96-NR; HR, 1.93; CI95, 0.58-6.47; p-value=.29) (Figure 4B).

**Figure 4:**
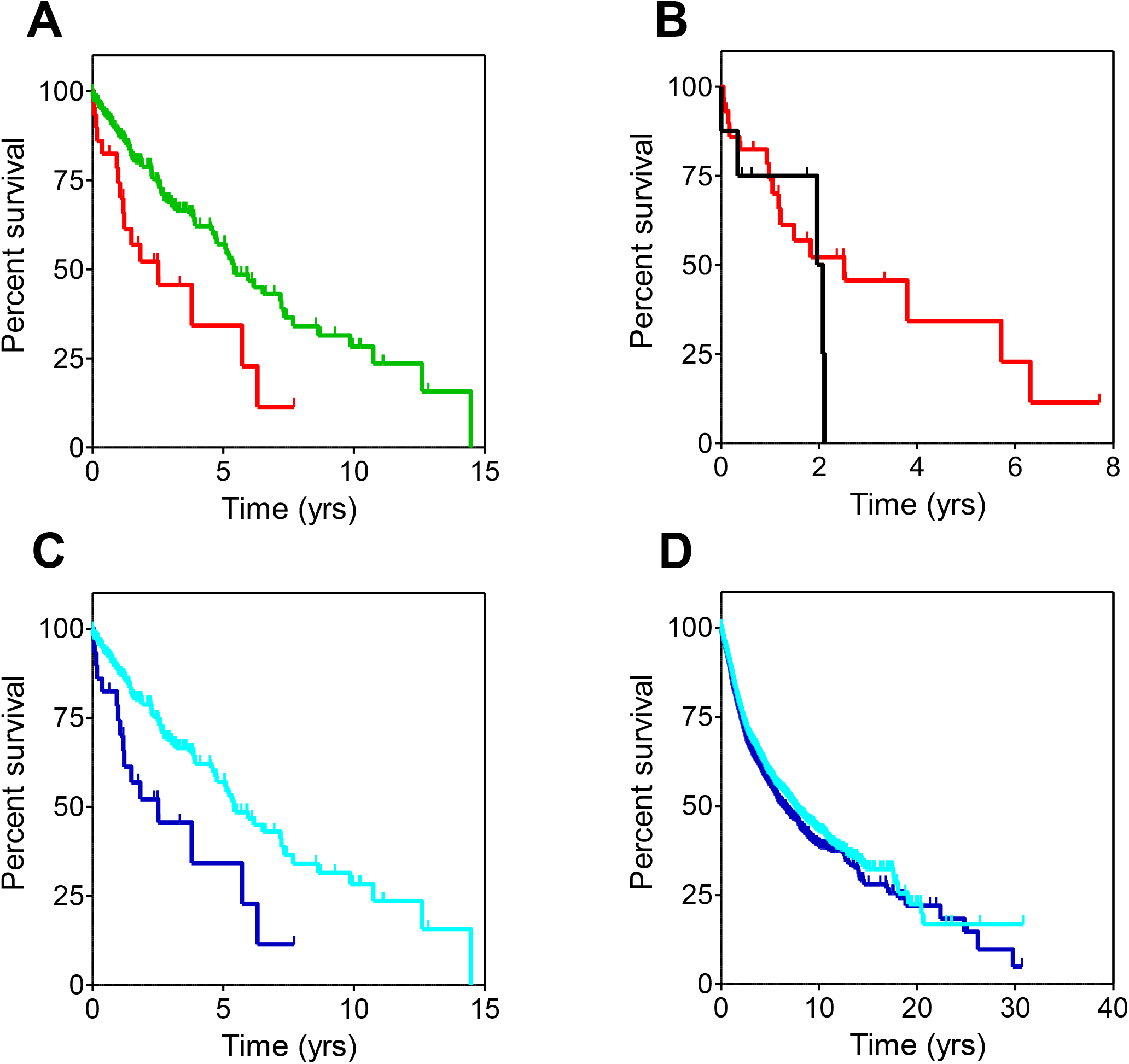
Overall survival curves from the study of eNSCLC patients. **A**: stage I eNSCLC patients sorted by cluster. **B**: stage I eNSCLC patients belonging to cluster 0 and stage IV eNSCLC patients. **C**: stage I eNSCLC patients sorted by the presence or absence of mutations in any of the 167 genes described in the text. **D**: all patients included in the TCGA dataset sorted by the presence or absence of mutations in any of the above genes. Censored patients are denoted with spikes. Color codes are the same for all panels: red for cluster 0, green for cluster 1, black for stage IV eNSCLC, dark blue for patients with mutations and light blue for patients without mutations in the intended genes.

As in the previous CRC case, analysis of the mutational burden showed that patients belonging to cluster 0 showed mutations in 3,325 genes, whilst those belonging to cluster 1 showed mutations in 16,541 genes. 3,158 of these genes were mutated in both clusters, therefore 167 genes (shown in Suppl. Table 5) were mutated exclusively in stage I eNSCLC patients assigned to cluster 0 and might be responsible for their worse prognosis, hence suggesting those gene mutations could be used as surrogate markers for cluster 0. Indeed, as seen in Fig. 4C, the MOS of patients harboring mutations in these genes (n=29; MOS, 2.5 years; CI95, 1.2-NR) was significantly worse than that of patients with no mutations (n=237; MOS, 5.4 years; CI95, 4.8-7.3; HR, 2.22; CI95, 1.32-3.85; p-value=.003). Moreover, when mutations in these 167 genes were evaluated along the entire TCGA dataset regardless their cancer origin (Fig. 4D), patients harboring mutations (n=3,777) had a slightly worse survival than those not harboring them (n=7,907). MOS were 6.3 years (CI95, 5.71-7.09) and 7.5 years (CI95, 6.98-7.97) for patients with mutations and without mutations, respectively (HR, 1.10; CI95, 1.05-1.20; p-value=.001). Therefore, the presence of mutations in these 167 genes seemed to be an indicator of negative prognosis for most cancer patients, particularly for eNSCLC patients diagnosed at stage I. As stated above for the colorectal cancer study, more work is warranted to ascertain their suitability as biomarkers of negative prognosis as well as their potential as novel therapeutic targets.

### Pheochromocytoma

Differences in OS were observed between patients from the adrenal cancer group depending on their inclusion on cluster 0 and 1, with the former showing better prognosis than the latter (Fig. 2C). When segregating the different pathologies included in the group (described in Suppl. Table 3), only pheochromocytoma (PHEO) patients exhibited differences in the survival pattern between the two clusters (Fig. 5A). Notably, the study was affected by an unusually high number of censored patients, so these results must be handled with caution. Likewise, the longer OS precluded from obtaining MOS values and made statistical significance rather borderline, but still patients in cluster 1 (n=21) showed an increased risk of death with respect to those in cluster 0 (n=126) (HR, 7.21; CI95, 1.01-51.48; p-value=.049).

**Figure 5:**
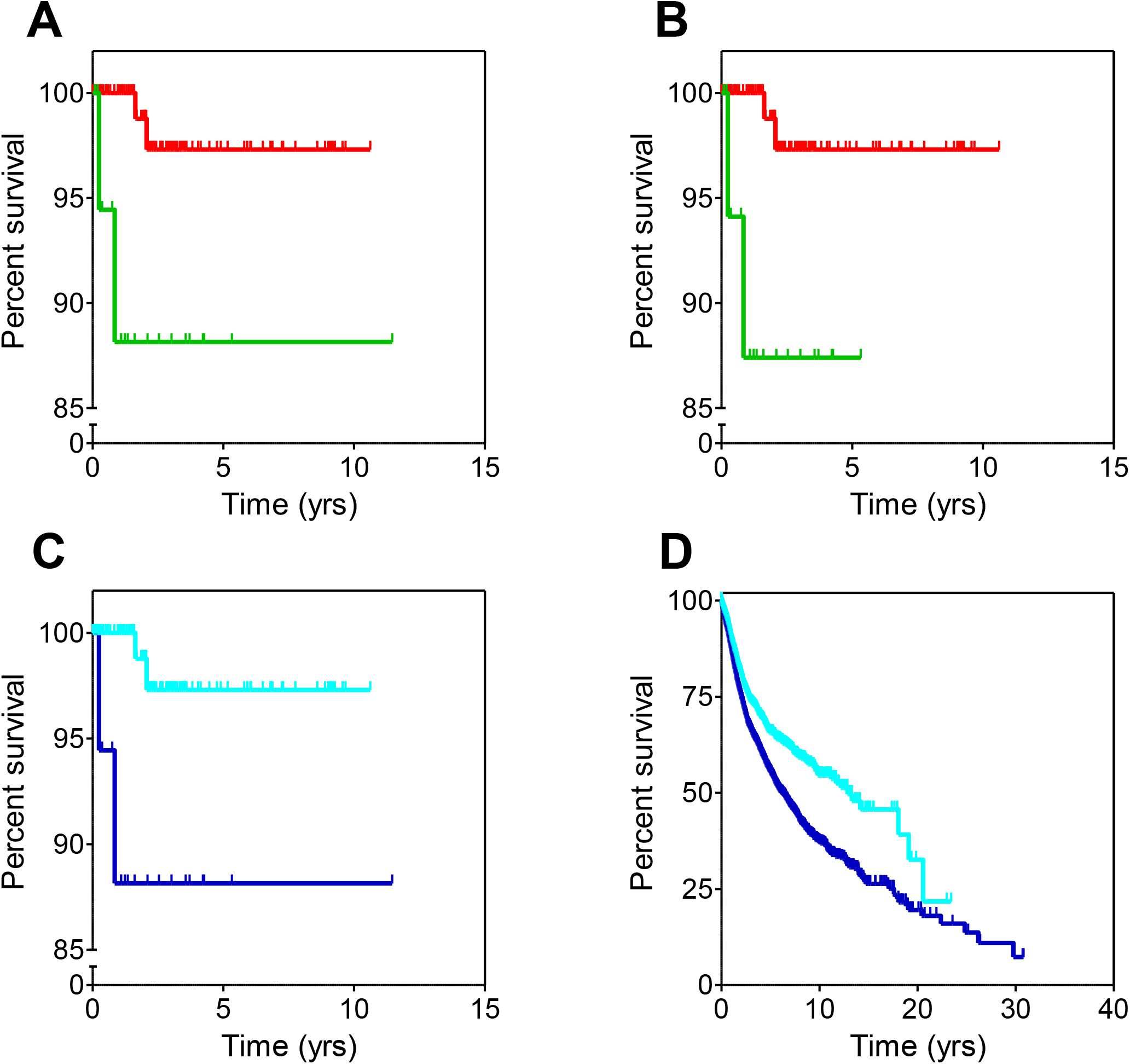
Overall survival curves from the study of pheochromocytoma (PHEO) patients. **A**: all PHEO patients sorted by cluster. **B**: non-metastatic PHEO patients sorted by cluster. **C**: non-metastatic PHEO patients sorted by the presence or absence of mutations in any of the 313 genes described in the text. **D**: all patients included in the TCGA dataset sorted by the presence or absence of mutations in any of the above genes. Censored patients are denoted with spikes. Color codes are the same for all panels: red for cluster 0, green for cluster 1, dark blue for patients with mutations and light blue for patients without mutations in the intended genes.

Stratification analysis showed that cluster assignment only influenced the survival of non-metastatic PHEO patients. Those belonging to cluster 0 (n=126) had better OS than those belonging to cluster 1 (n=20) (Figure 5B; HR, 7.89; CI95, 1.10-56.46; p-value=.04). As above, the unusually high censoring urges to consider these results cautiously. Analysis of the mutational burden revealed that PHEO cluster 0 patients carried mutations in 1,034 genes whilst PHEO cluster 1 patients did so in 359 genes, 46 of these genes being common to both clusters. Therefore, 313 genes (listed in Suppl. Table 6) were exclusively mutated in patients from cluster 1, the one with worse prognosis.

We then evaluated whether those gene mutations could be used as surrogate markers for cluster 1. Although MOS values were not reached, Fig. 5C shows that non-metastatic PHEO patients with mutations in any of the 313 genes (n=21) were at significantly higher risk of death than those presenting no mutations (n=126) (HR, 7.14; CI95, 1.01-50.0; p-value=.049). A similar analysis was performed in the TCGA dataset: survival analysis was performed dividing these patients in two groups depending on whether they did or did not harbor mutations in any of the 313 genes (Figure 5D). Interestingly, patients with mutations (n=8,933) showed significantly worse survival curves than patients in whom these genes remained unmutated (n=2,751). MOS were 6.1 years (CI95, 5.71-6.62) and 12.85 (CI95, 11.18-NR) for patients with mutated genes and patients without mutations, respectively (HR, 1.45; CI95, 1.33-1.59; p-value <.0001).

The results obtained with these genes in the whole set of cancer patients included in the TCGA dataset suggest that the scrutiny of the data from PHEO patients, despite the limitations caused by their high censoring, has allowed the identification of 313 genes that may be responsible for poorer survival in cancer. As in previous cases, more work should be done to explore in more detail the role of these genes in cancer and their potential value as prognostic biomarkers or even as possible new therapeutic targets.

### Urothelial carcinoma

Patients included in the here called urinary tract cancer group belonging to cluster 0 exhibited worse survival curves (Fig. 2D) with statistically significant shorter MOS than those included in cluster 1. Most of these patients were diagnosed with urothelial carcinoma, and differences in survival remained for them as displayed in Figure 6A. MOS were 1.5 years (CI95, 0.97-NR) and 2.9 years (CI95, 2.13-5.40) for urothelial carcinoma patients belonging to cluster 0 (n=29) and cluster 1 (n=368), respectively (HR, 1.82; CI95, 1.11-3.03; p-value=.018). As in previous cases, analysis of the mutational burden showed that patients in cluster 0 carried mutations in 2,428 genes whilst patients in cluster 1 showed mutations in 17,573 genes. 2,376 of these mutated genes were common to both clusters, hence 52 genes were exclusively mutated in patients from cluster 0 and might be responsible for their worse survival. No significant differences were observed in OS when selecting patients with mutations only in these 52 genes (data not shown), indicating that these could not be used as surrogate biomarkers for cluster 0. Hence this analysis was not pursued further.

**Figure 6:**
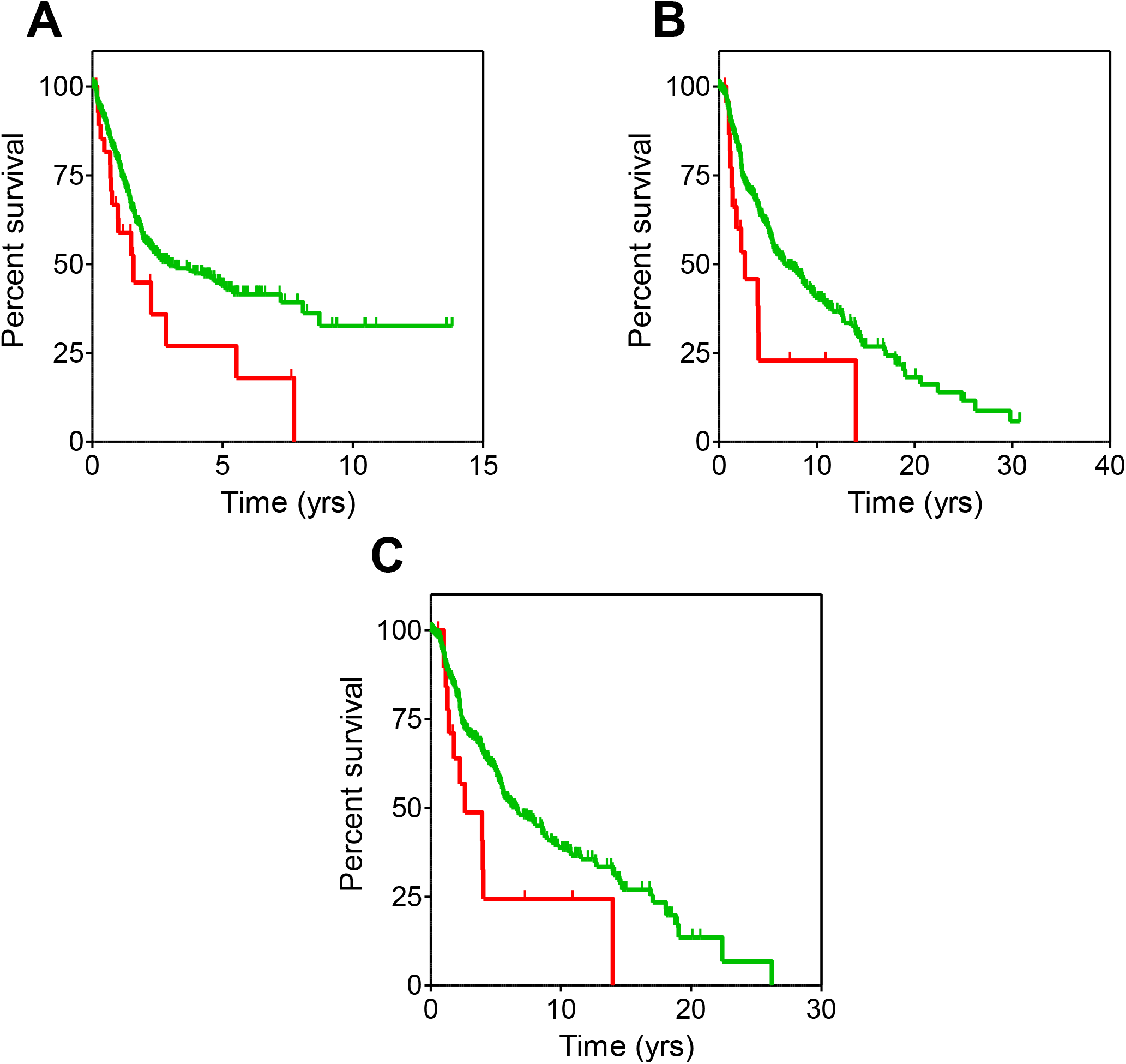
Survival curves from the study of urothelial carcinoma and melanoma patients. **A**: urothelial carcinoma patients sorted by cluster. **B**: melanoma patients sorted by cluster. **C**: non-metastatic melanoma patients sorted by cluster. Censored patients are denoted with spikes. Color codes are the same for all panels: red for cluster 0, green for cluster 1.

### Melanoma

Although no differences in survival were observed between the two clusters in the skin cancer group, the situation changed when the group was divided in its two major disease subsets, melanoma, and ocular melanoma. Only in the former case, patients included in cluster 0 (n=23) showed significantly worse OS than those in cluster 1 (n=429) (Figure 6B). MOS were 2.6 years (CI95, 1.39-NR) and 6.7 years (CI95, 5.54-8.92) for cluster 0 and cluster 1 patients, respectively (HR, 2.17; CI95, 1.33-3.70; p-value=.005). Stratification analysis showed that cluster assignment only influenced the survival of non-metastatic melanoma patients. As observed in Fig. 6C, those belonging to cluster 0 (n=20) had a higher risk of death (HR, 1.96; CI95, 1.09-3.57; p-value=.025) than those belonging to cluster 1 (n=368). Median OS were 2.6 years (CI95, 1.78-NR) and 6.5 years (CI95, 5.49-8.60) for patients in cluster 0 and cluster 1, respectively. Analysis of the mutational burden showed that it was significantly higher for patients in cluster 1 (17,665 genes) than for the worse prognosis cluster 0 patients (815 genes). Interestingly, as many as 814 genes were common to both clusters, therefore one single gene was exclusively mutated in melanoma cluster 0 patients: SHARPIN, a regulator of melanomagenesis already described to promote melanoma proliferation, metastasis, and invasion [27].

### Validation with a different dataset: breast cancer

To further validate GFPrint^™^, a different dataset was used with the goal of demonstrating that the performance of the tool was similar to that observed with the TCGA dataset and reliable results could be obtained. In this case, WES and clinical information from 945 breast tumors were obtained from Broad Institute of MIT and Harvard. As above, sequences were encrypted and processed with GFPrint^™^ to distribute the patients in the same latent space previously generated (Fig. 7A) and 2 subpopulations of patients were now identified by the tool: SP0 (n=358) and SP1 (n=587). These subpopulations yielded significant differences in their survival progression at 7 years (Fig. 7B), with patients in SP1 being at a higher risk of death compared to those in SP0 (HR, 1.48; CI95, 1.01-2.16; p-value=.046).

**Figure 7:**
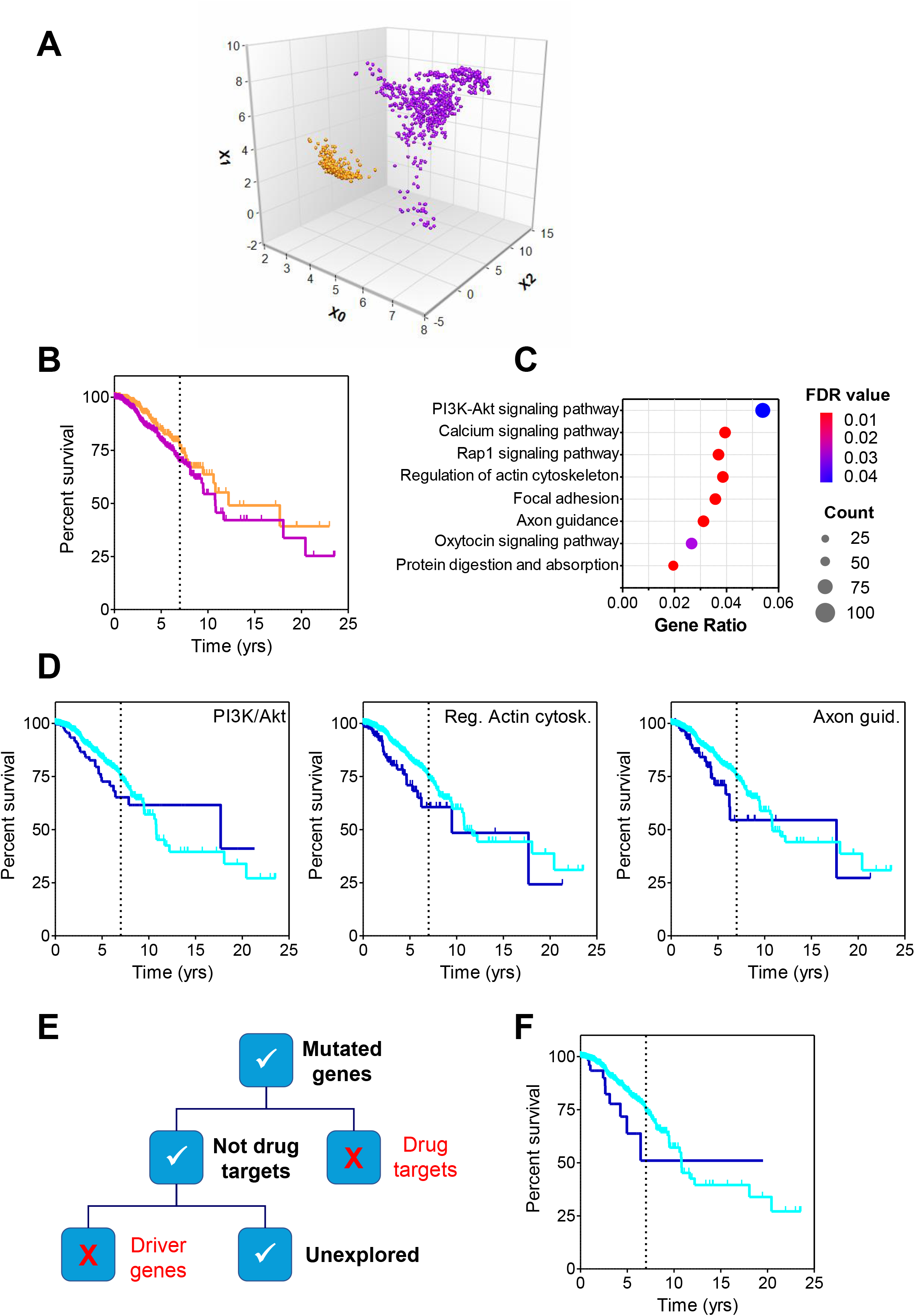
Study of breast cancer dataset from Broad Institute of MIT and Harvard. **A**: Latent space generated by GFPrint^™^ for the dataset; two subpopulations of patients were identified, SP0 (N = 358) denoted in orange, and SP1 (N = 587) in purple. **B**: Survival curves for patients in SP0 and SP1 (same colors as above). The vertical dotted line highlights the 7-year time point. **C**: Outcome of the PEA run using KEGG. The eight most significantly (FDR < 0.05) enhanced pathways are displayed in the vertical axis, whilst the horizontal axis shows the gene ratio (i.e., the number of analyzed genes that belong to a given pathway divided by the total number of genes included in that pathway). As explained in the legend, dots sizes are proportional to the number of analyzed genes present in that pathway with colors corresponding to the FDR value. **D**: Survival curves for patients included in the dataset sorted by the presence (dark blue) or absence (light blue) of mutations in genes belonging to the PI3K/Akt signaling, regulation of actin cytoskeleton and axon guidance pathways as labeled. The vertical dotted line in each graph highlights the 7-year time point. **E**: Diagram showing the criteria followed for triaging the mutated genes included in the three pathways above with the aim of selecting novel genes not previously associated with cancer. **F**: Survival curves for patients included in the dataset sorted by the presence (dark blue) or absence (light blue) of mutations in the 75 genes belonging to the PI3K/Akt signaling pathway that were selected after the triage. The vertical dotted line highlights the 7-year time point.

Mutational burden analysis showed that the number of mutated genes in SP1 patients was higher than in SP0 ones (5,520 and 1,491 genes respectively, with 692 of them being common to both groups). To gain insight on the genes that may influence the worse evolution of SP1 patients, PEA was performed to identify biological pathways and functions that were significantly overrepresented (more than expected by chance) in the SP1 mutated genes. In an effort to make it comprehensive, the analysis was performed using several databases, namely KEGG, Wikipathways and Gene Ontology. The most compelling results were obtained with KEGG, and they are shown in Figure 7C: eight different pathways were identified with an adjusted p-value (FDR) below .05. For each of these pathways, differences in survival were investigated among all patients included in the dataset according to whether or not they carried mutations in any of the genes included in that pathway. Only three of them (PI3K/Akt signaling, 130 genes; regulation of actin cytoskeleton, 93 genes; and regulation of axon guidance, 75 genes) showed a higher risk of death at 7 years for patients harboring mutations in any of their genes (Fig 7D). Patients with mutations in genes belonging to PI3K/Akt pathway (n=206), actin cytoskeleton pathway (n=128) and axon guidance pathway (n=127) showed an increased risk of death of 1.6 (CI95, 1.07-2.48; p-value=.023), 2.1 (CI95, 1.38-3.41; p-value=.001) and 1.8 (CI95, 1.16-3.08; p-value=.01), respectively.

The mutated genes included in these three pathways were triaged according to a list of criteria aimed at selecting novel genes not previously associated with cancer (Fig. 7E). First, genes whose products were known to be cancer drug targets were removed. Second, those deemed as cancer driver genes were also discarded. This left only 75, 54 and 45 genes from the PI3K/Akt signaling, regulation of actin cytoskeleton and axon guidance pathways, respectively. As above, differences in survival were investigated among the patients carrying or not mutations in these selected genes: only the 75 genes selected from the PI3K/Akt signaling pathway (shown in Suppl. Table 7) were related to a higher risk of death at 7 years (HR, 2.26; CI95, 1.17-4.36; p-value=.016) (Figure 7F). It is therefore tempting to hypothesize that mutations on these 75 genes that had not been considered previously in oncology may be influencing a worse evolution of the disease in breast cancer patients.

To gain more information on the nature of these 75 genes, originally selected from the PEA run in KEGG, a new PEA was performed on them using Reactome [28] as the source database: this should render a transversal insight from both databases that may enrich the ultimate outcome. Analysis showed that 21 pathways were significantly enriched in these 75 genes (FDR < .01). Interestingly, 8 of the 10 most significant ones (FDR < 1E-4) belonged to the hierarchy of extracellular matrix organization (ECM), including the 6 top significant pathways: laminin interactions (FDR 5.41E-14; 18 identified genes of the 31 in the pathway), ECM proteoglycans (FDR 5.41E-14; 26 of 79), anchoring fibril formation (FDR 1.87E-10, 8 of 15), non-integrin membrane-ECM interactions (FDR 5.66E-10; 19 of 61), assembly of collagen fibrils and other multimeric structures (FDR 7.60E-10; 13 of 67) and collagen chain trimerization (FDR 5.65E-7; 10 of 44). Given the essential role of ECM remodeling in tumor metastasis, it might not be unreasonable to conclude that a major propensity to metastasis may underlie the poorer prognosis of the breast cancer patients included in this subpopulation. Likewise, as in some cases described above, it is tempting to think of these genes as a suitable panel to identify patients at risk, regardless of their possible role as therapeutic targets. Again, experimental work should be done to demonstrate the validity of these hypothesis.

## DISCUSSION

In this study, we have used GFPrint^™^ to transform high-dimensional, noisy gene sequencing data to a lower dimensional, meaningful representation. This was used to find patterns in genomic data to identify patient subpopulations with a particular clinical outcome. First, we developed GFPrint^™^, a proprietary streaming algorithm to deeply extract functional features from high dimensional gene sequencing analysis. Next, we examined the extracted embedding representation with different unsupervised models to determine nonlinear relationships in the data to detect different clusters of patients that would behave with a different clinical outcome. Lastly, we analyzed the lower-dimensional representations by mapping back to the original data to identify the aggregated mutational burden on genes and ablated biological pathways that could play critical roles and serve as virtual biomarkers for cancer prognostics and target identification. This knowledge would also trigger new hypotheses for cancer survival analysis in a genomic cancer research setting. GFPrint^™^ presents many advantages: (i) it works in small or large sequencing datasets; (ii) it can learn from millions of sequences and transfer the learnt knowledge to unseen datasets; (iii) it can tolerate a moderate degree of noise and mislabeled training data; (iv) it can generate embedding spaces fully automatically, alleviating the need for careful and time-consuming hand-tuning; and (v) it can be accessed through a secured web portal system that allow researchers to upload their existing anonymized genetic data.

The profound advances observed in the next generation sequencing arena over the last decade are revolutionizing the field of genomics and more particularly its application to medicine. Novel instruments and technologies allow the parallel sequencing of millions to billions of DNA and RNA fragments, making the sequencing process highly efficient and reliable. These technological advancements are concomitant to a substantial decrease in the costs associated to their use. Therefore, it is foreseeable that genome sequencing of samples from patients will become a routine habit in clinical settings in the not-too-distant future. The greater amount of high dimensional genomic data will allow to discover more molecular biomarkers in different clinical fields, for instance, cancer survival and or treatment. The flipside of these developments is that having all this information at the clinicians and investigators disposal may be in vain if appropriate tools to exploit and interpret it are not available. It follows that the accessibility to this kind of tools is an urgent need. Machine learning methods can capitalize on this information, as they have the capability of finding unpredictable, often nonlinear, interactions between somatic point mutations and different clinical endpoints, which may contribute to more accurate prognostic or predictive medicine. In relation with the abovementioned problem, this is commonly approached with transfer-learning, where a model which has been trained on a single task (e.g., genetics of abundant cancers) is used to fine-tune on an unrelated target task (e.g. cancer survival).

Precisely, GFPrint^™^ has been designed with the goal of contributing to fill this unmet need and the results presented here confirm that the tool is well suited for that purpose. By analyzing large cancer genomic datasets, GFPrint^™^ can generate virtual representations of tumor genetics regardless their tissue of origin, distributed throughout latent spaces where it is possible to identify subpopulations that show different clinical outcomes. Since these subpopulations are defined by their position in the latent space, and because these positions are dictated by their genetic profile, it follows that the observed different clinical outcomes must be strongly influenced by the genetic features of the patients, more particularly their mutational landscape. For instance, in stage I eNSCLC patients, GFPrint^™^ identified 167 mutated genes significantly correlated with a higher risk of death demonstrating that it can effectively leverage high-dimensional genetic data that is relevant but not known to be related to the problem of interest, in this case, cancer survival. Independently from their site of origin, the tool was able to learn from scratch with a significantly large number of samples while implicitly associating and prioritizing tumor genotypes based on their contribution to survival. This valuable information can be further exploited by delving such mutational landscape in search of genes that can be harnessed for many different purposes such as cancer prognosis or new target identification. Indeed, GFPrint^™^ is potentially applicable to many complex diseases, and our overall approach is expected to be valuable in developing clinical tests that incorporate personal genome sequences into disease-risk prediction.

More specifically, the analysis performed with the TCGA dataset has led to the identification of panels of genes whose mutations seem to dictate the overall survival of non-metastatic patients with colorectal cancer, epidermoid non-small cell lung cancer or pheochromocytoma whose prognosis turned out to be worse than expected based on their non-metastatic status. This was particularly evident for stage I eNSCLC patients: the subgroup harboring mutations in a panel of 167 genes showed disease progression as bad as that of stage IV patients and, therefore, this panel of genes appears as a biomarker of worse prognosis with greater predictive value than disease stage. The same applies to CRC and PHEO patients, with the 15 and 313 genes respectively identified showing stronger predictive value than metastatic state of the patients. Applying this procedure to melanoma patients has unveiled SHARPIN, a gene known to be involved in melanomagenesis and disease progression, as being critical for the worse survival of patients, a result that confirms the performance of the tool.

The results obtained with the breast cancer dataset validate GFPrint^™^ further. The latent space that was generated with the wider TCGA dataset was used to arrange the virtual representations created by the tool for the patients included in this distinct dataset, and still GFPrint^™^ was able to discern two unique subpopulations with differentiated clinical outcomes that could be investigated, leading to interesting conclusions. It is noteworthy that most of the PI3K/Akt signaling route genes whose mutations dictated worse survivals were also involved in cellular pathways related to the extracellular matrix reorganization, a process widely recognized to be related to metastasis as thoroughly reviewed in the bibliography [29-31]. Therefore, the conclusion of the analysis is not only reasonable and consistent (hence endorsing the performance of GFPrint^™^) but may also provide valuable clues for identifying novel points of therapeutic intervention, whether they are independent targets or part of mutually dependent groups of genes underlying synthetic lethality opportunities [32] to be exploited in novel, selective and efficient treatments.

With all of the above, it is clear that GFPrint^™^ has value as a hypothesis generation tool of broad application for genome-informed therapies. Besides providing rationales to propel new investigations intended to understand and treat diseases more efficiently, the tool can help designing personalized therapies tailored to treat groups of patients with a common genetic signature encompassing different conditions, a complementary approach to the classical indication-centric treatments. At the beginning of 2024, only 7 tissue-agnostic cancer treatments had been approved by the FDA [33], a number expected to increase soon. This, in turn, must contribute to improving the poor success rates of current genome-informed therapies in oncology, where the response in the USA was estimated to be only 11% in 2020, a figure that may be influenced by the low eligibility of patients, reaching only 13% [34]. As previously stated, the lack of appropriate tools to exploit the increasingly available genomic data from patients is one of the reasons explaining this unsatisfactory landscape, and tools like GFPrint^™^ should contribute to improve the situation.

A recent study [35] has been published showing the value of bioinformatics analysis for integrating genomic and clinical data and its relevance for precision medicine. Additionally, different publications have described the design and use of ML engines to exploit genetic information for the same purpose [36-38]. The latter tools condense the genetic information to be processed by the ML framework, e.g. by scoring individual mutations depending on how they affect the function of the encoded protein to ultimately generate aggregated scores. On the opposite, the encryption process at the helm of GFPrint^™^ preserves the granularity of the information available in the entire DNA sequencing, making it highly sensitive to differential features for each individual patient that may be lost through condensation. Likewise, such tools are based on the function of the proteins encoded by the mutated genes and are therefore restricted to the exomic DNA. However, GFPrint^™^ encryption system utilizes the whole genome sequence, therefore it is not restricted to specific genes. Additionally, those tools are dependent of external information related to the gain or loss of function of the mutated protein which is brought in during the embedding process, and such external information may introduce unwanted biases. Conversely, the embedding process in GFPrint^™^ is exclusively based on the naked DNA sequence and is therefore free from to external influences that may cause distortions. Finally, existing tools face scalability challenges, as their latent spaces are fixed and cannot incorporate new samples for comparison with the initial cohort. GFPrint^™^, however, offers scalability and flexibility, enabling the integration of new samples into established latent spaces for ongoing comparative analysis with a broader population.

Our work has several limitations. First, the TCGA dataset presented an unusual high level of censoring data. More than 75% of the patients in the dataset are censored. This urges to consider the survival results very cautiously. Second, although the most common tumor types are included in the dataset, their distribution does not match the relative population incidences (e.g., like those described in [38]), largely owing to differences among contributing research groups in the numbers of sequenced samples. Third, our analysis did not consider that each data acquisition technology has its own artifacts, biases, and limitations. We cannot exclude that the obtained results could reflect the different specificities of these unwanted effects in a large dataset as TCGA. However, TCGA data is a benchmark for DNA sequencing data, used in many different published works even without correcting for technology-specific biases and thus, we can assume that this is not a relevant matter. Four, GFPrint^™^ was tested on a large dataset of somatic point mutations with known cancer types (i.e., the histological biopsy sites were already known). Therefore, it only demonstrates the power of capturing complex association between somatic point mutations and biopsy confirmed cancer types. Whether this information could also be applied to tumor samples with completely unknown cancer type information (such as liquid biopsy data) is not known. Moreover, the association between point mutations and other genetic aberrations such as copy number variance, translocation, and small insertion and deletion is not covered by GFPrint^™^. However, our work shows that, for some tumor types, analyzing point mutations alone is good enough for retrieving those complex genetic associations correlated with a high risk of death. Five, we have found interesting molecular signatures for some tumors but not others. Whether this was related to the molecular biology of each disease or was due to all the previous caveats mentioned above is not clear. In this case, dedicated studies will be needed to clarify this subject. Finally, the system was validated only in one external dataset (from the Broad Institute). At any rate, our results raised potentially fruitful avenues for future translational research in cancer.

In summary, we demonstrate that GFPrint^™^ is an effective framework to leverage high-dimensional data that is relevant but not directly related to the problem of interest. In this study, the tool was able to extract meaningful features from gene sequencing data that were able to identify genetic signatures correlated to cancer survival. Although the studies shown in this paper have focused exclusively on cancer patients, GFPrint^™^ is a disease-agnostic tool of broad application in any therapeutic area where the genetic profile of patients influences the onset and evolution of the pathology. Furthermore, it may be applied to evaluate the genome of healthy individuals with the aim of identifying hidden genetic patterns and associations that may predispose to certain conditions. In overall, we hope that the versatility of the tool will help the scientific community to foster the development of a better and wiser medicine tailored to the idiosyncrasy and individual circumstances of each patient.

## DATA AVAILABILITY

The datasets analyzed during the current study are available in the Science Data Bank repository, https://doi.org/10.57760/sciencedb.17342

They were originally obtained from the following sites: TCGA pan-cancer database, available at https://gdc.cancer.gov/

Broad Institute of MIT and Harvard dataset available at http://firebrowse.org/?cohort=BRCA

## ACKNOWLEDGEMENTS

This work was partly funded by CDTI (Centro para el Desarrollo Tecnológico e Industrial), through the Neotec Project SNEO-20211030.

The results presented here are in part based upon data generated by the TCGA Research Network at https://www.cancer.gov/tcga.

## AUTHOR CONTRIBUTIONS

Conceptualization, JMD

Data curation, GSM, DPM

Formal analysis, GSM, PGC, JCI

Software, GSM, PGC, JCI

Supervision, JMD

Writing – original draft, GSD, JMD

Writing – review & editing, DPM, JMD

