## Supplementary Tables for "GFPrint™: A MACHINE LEARNING TOOL FOR TRANSFORMING GENETIC DATA INTO CLINICAL INSIGHTS"

**SUPPLEMENTARY TABLES – G. Sanz-Martín *et al*.**

**Supplementary Table 1: Distribution of the 145 tumor histotypes found in the TCGA dataset along the 25 cancer groups created according to the primary diagnosis described there.**

NOS, not otherwise specified.

| Cancer group | Histotype | Primary diagnosis |
|---|---|---|
| Adrenal | Adrenal gland, NOS | Ganglioneuroblastoma |
| | Adrenal gland, NOS | Neuroblastoma, NOS |
| | Adrenal gland, NOS | Paraganglioma, malignant |
| | Adrenal gland, NOS | Pheochromocytoma, malignant |
| | Adrenal gland, NOS | Pheochromocytoma, NOS |
| | Cortex of adrenal gland | Adrenal cortical carcinoma |
| | Cortex of adrenal gland | Ganglioneuroblastoma |
| | Cortex of adrenal gland | Neuroblastoma, NOS |
| | Cortex of adrenal gland | Pheochromocytoma, NOS |
| | Medulla of adrenal gland | Neuroblastoma, NOS |
| | Medulla of adrenal gland | Pheochromocytoma, malignant |
| | Retroperitoneum | Paraganglioma, NOS |
| | Retroperitoneum | Pheochromocytoma, NOS |
| Bone | Bone, NOS | Ewing sarcoma |
| | Bone, NOS | Not Reported |
| | Bone, NOS | Undifferentiated sarcoma |
| | Short bones of lower limb and associated joints | Leiomyosarcoma, NOS |
| Breast | Breast, NOS | Apocrine adenocarcinoma |
| | Breast, NOS | Cribriform carcinoma, NOS |
| | Breast, NOS | Infiltrating duct and lobular carcinoma |
| | Breast, NOS | Infiltrating duct carcinoma, NOS |
| | Breast, NOS | Infiltrating duct mixed with other types of carcinoma |
| | Breast, NOS | Infiltrating lobular mixed with other types of carcinoma |
| | Breast, NOS | Intraductal micropapillary carcinoma |
| | Breast, NOS | Intraductal papillary adenocarcinoma with invasion |
| | Breast, NOS | Large cell neuroendocrine carcinoma |
| | Breast, NOS | Lobular carcinoma, NOS |
| | Breast, NOS | Medullary carcinoma, NOS |
| | Breast, NOS | Metaplastic carcinoma, NOS |
| | Breast, NOS | Mucinous adenocarcinoma |
| | Breast, NOS | Paget disease and infiltrating duct carcinoma of breast |
| | Breast, NOS | Phyllodes tumor, malignant |
| | Breast, NOS | Tubular adenocarcinoma |
| | Lower-inner quadrant of breast | Infiltrating duct carcinoma, NOS |

| Cancer group | Histotype | Primary diagnosis |
| --- | --- | --- |
| | Lower-inner quadrant of breast | Infiltrating duct mixed with other types of carcinoma |
| | Lower-outer quadrant of breast | Infiltrating duct carcinoma, NOS |
| | Overlapping lesion of breast | Infiltrating duct carcinoma, NOS |
| | Overlapping lesion of breast | Pleomorphic carcinoma |
| | Upper-inner quadrant of breast | Infiltrating duct carcinoma, NOS |
| | Upper-inner quadrant of breast | Infiltrating lobular mixed with other types of carcinoma |
| | Upper-outer quadrant of breast | Infiltrating duct carcinoma, NOS |
| CNS | Brain, NOS | Astrocytoma, anaplastic |
| | Brain, NOS | Astrocytoma, NOS |
| | Brain, NOS | Glioblastoma |
| | Brain, NOS | Gliosarcoma |
| | Brain, NOS | Mixed glioma |
| | Brain, NOS | Oligodendroglioma, anaplastic |
| | Brain, NOS | Oligodendroglioma, NOS |
| | Cerebrum | Astrocytoma, anaplastic |
| | Cerebrum | Astrocytoma, NOS |
| | Cerebrum | Mixed glioma |
| | Cerebrum | Oligodendroglioma, anaplastic |
| | Cerebrum | Oligodendroglioma, NOS |
| | Frontal lobe | Astrocytoma, anaplastic |
| | Frontal lobe | Astrocytoma, NOS |
| | Frontal lobe | Glioblastoma |
| | Frontal lobe | Mixed glioma |
| | Frontal lobe | Oligodendroglioma, anaplastic |
| | Occipital lobe | Astrocytoma, anaplastic |
| | Occipital lobe | Glioblastoma |
| | Overlapping lesion of brain | Glioblastoma |
| | Parietal lobe | Glioblastoma |
| | Parietal lobe | Mixed glioma |
| | Temporal lobe | Astrocytoma, anaplastic |
| | Temporal lobe | Glioblastoma |
| | Temporal lobe | Mixed glioma |
| | Temporal lobe | Oligodendroglioma, NOS |
| Connective and soft tissue | Connective, subcutaneous and other soft tissues of abdomen | Adenocarcinoma, NOS |
| | Connective, subcutaneous and other soft tissues of abdomen | Extra-adrenal paraganglioma, NOS |
| | Connective, subcutaneous and other soft tissues of abdomen | Leiomyosarcoma, NOS |
| | Connective, subcutaneous and other soft tissues of head, face, and neck | Malignant peripheral nerve sheath tumor |
| | Connective, subcutaneous and other soft tissues of head, face, and neck | Pleomorphic liposarcoma |
| | Connective, subcutaneous and other soft tissues of head, face, and neck | Synovial sarcoma, spindle cell |
| | Connective, subcutaneous and other soft tissues of head, face, and neck | Undifferentiated sarcoma |

| Cancer group | Histotype | Primary diagnosis |
|---|---|---|
| | Connective, subcutaneous and other soft tissues of lower limb and hip | Dedifferentiated liposarcoma |
| | Connective, subcutaneous and other soft tissues of lower limb and hip | Fibromyxosarcoma |
| | Connective, subcutaneous and other soft tissues of lower limb and hip | Giant cell sarcoma |
| | Connective, subcutaneous and other soft tissues of lower limb and hip | Leiomyosarcoma, NOS |
| | Connective, subcutaneous and other soft tissues of lower limb and hip | Malignant fibrous histiocytoma |
| | Connective, subcutaneous and other soft tissues of lower limb and hip | Malignant peripheral nerve sheath tumor |
| | Connective, subcutaneous and other soft tissues of lower limb and hip | Synovial sarcoma, biphasic |
| | Connective, subcutaneous and other soft tissues of lower limb and hip | Synovial sarcoma, spindle cell |
| | Connective, subcutaneous and other soft tissues of lower limb and hip | Undifferentiated sarcoma |
| | Connective, subcutaneous and other soft tissues of pelvis | Dedifferentiated liposarcoma |
| | Connective, subcutaneous and other soft tissues of pelvis | Extra-adrenal paraganglioma, NOS |
| | Connective, subcutaneous and other soft tissues of pelvis | Fibromyxosarcoma |
| | Connective, subcutaneous and other soft tissues of pelvis | Leiomyosarcoma, NOS |
| | Connective, subcutaneous and other soft tissues of pelvis | Malignant fibrous histiocytoma |
| | Connective, subcutaneous and other soft tissues of pelvis | Undifferentiated sarcoma |
| | Connective, subcutaneous and other soft tissues of thorax | Aggressive fibromatosis |
| | Connective, subcutaneous and other soft tissues of thorax | Dedifferentiated liposarcoma |
| | Connective, subcutaneous and other soft tissues of thorax | Extra-adrenal paraganglioma, NOS |
| | Connective, subcutaneous and other soft tissues of thorax | Fibromyxosarcoma |
| | Connective, subcutaneous and other soft tissues of thorax | Leiomyosarcoma, NOS |
| | Connective, subcutaneous and other soft tissues of thorax | Malignant fibrous histiocytoma |
| | Connective, subcutaneous and other soft tissues of thorax | Malignant peripheral nerve sheath tumor |
| | Connective, subcutaneous and other soft tissues of thorax | Paraganglioma, NOS |
| | Connective, subcutaneous and other soft tissues of thorax | Synovial sarcoma, NOS |
| | Connective, subcutaneous and other soft tissues of thorax | Synovial sarcoma, spindle cell |
| | Connective, subcutaneous and other soft tissues of thorax | Undifferentiated sarcoma |
| | Connective, subcutaneous and other soft tissues of trunk, NOS | Extra-adrenal paraganglioma, NOS |
| | Connective, subcutaneous and other soft tissues of trunk, NOS | Fibromyxosarcoma |
| | Connective, subcutaneous and other soft tissues of trunk, NOS | Leiomyosarcoma, NOS |
| | Connective, subcutaneous and other soft tissues of trunk, NOS | Malignant fibrous histiocytoma |
| | Connective, subcutaneous and other soft tissues of trunk, NOS | Malignant peripheral nerve sheath tumor |

| Cancer group | Histotype | Primary diagnosis |
|---|---|---|
| | Connective, subcutaneous and other soft tissues of trunk, NOS | Synovial sarcoma, biphasic |
| | Connective, subcutaneous and other soft tissues of trunk, NOS | Undifferentiated sarcoma |
| | Connective, subcutaneous and other soft tissues of upper limb and shoulder | Dedifferentiated liposarcoma |
| | Connective, subcutaneous and other soft tissues of upper limb and shoulder | Leiomyosarcoma, NOS |
| | Connective, subcutaneous and other soft tissues of upper limb and shoulder | Synovial sarcoma, NOS |
| | Connective, subcutaneous and other soft tissues of upper limb and shoulder | Undifferentiated sarcoma |
| | Connective, subcutaneous and other soft tissues, NOS | Alveolar rhabdomyosarcoma |
| | Connective, subcutaneous and other soft tissues, NOS | Leiomyosarcoma, NOS |
| | Connective, subcutaneous and other soft tissues, NOS | Undifferentiated sarcoma |
| | Overlapping lesion of connective, subcutaneous and other soft tissues | Leiomyosarcoma, NOS |
| | Peripheral nerves and autonomic nervous system of upper limb and shoulder | Malignant peripheral nerve sheath tumor |
| | Retroperitoneum | Malignant peripheral nerve sheath tumor |
| | Spinal meninges | Malignant peripheral nerve sheath tumor |
| Esophagus | Esophagus, NOS | Adenocarcinoma, NOS |
| | Esophagus, NOS | Squamous cell carcinoma, keratinizing, NOS |
| | Esophagus, NOS | Squamous cell carcinoma, NOS |
| | Lower third of esophagus | Adenocarcinoma, NOS |
| | Lower third of esophagus | Basaloid squamous cell carcinoma |
| | Lower third of esophagus | Mucinous adenocarcinoma |
| | Lower third of esophagus | Squamous cell carcinoma, keratinizing, NOS |
| | Lower third of esophagus | Squamous cell carcinoma, NOS |
| | Lower third of esophagus | Tubular adenocarcinoma |
| | Middle third of esophagus | Adenocarcinoma, NOS |
| | Middle third of esophagus | Squamous cell carcinoma, NOS |
| | Thoracic esophagus | Adenocarcinoma, NOS |
| | Thoracic esophagus | Squamous cell carcinoma, keratinizing, NOS |
| | Upper third of esophagus | Squamous cell carcinoma, NOS |
| Head and neck | Anterior floor of mouth | Squamous cell carcinoma, keratinizing, NOS |
| | Anterior floor of mouth | Squamous cell carcinoma, NOS |
| | Base of tongue, NOS | Basaloid squamous cell carcinoma |
| | Base of tongue, NOS | Squamous cell carcinoma, keratinizing, NOS |
| | Base of tongue, NOS | Squamous cell carcinoma, large cell, nonkeratinizing, NOS |
| | Base of tongue, NOS | Squamous cell carcinoma, NOS |
| | Border of tongue | Squamous cell carcinoma, NOS |
| | Cheek mucosa | Basaloid squamous cell carcinoma |
| | Cheek mucosa | Squamous cell carcinoma, keratinizing, NOS |

| Cancer group | Histotype | Primary diagnosis |
|---|---|---|
| | Cheek mucosa | Squamous cell carcinoma, NOS |
| | Floor of mouth, NOS | Basaloid squamous cell carcinoma |
| | Floor of mouth, NOS | Squamous cell carcinoma, keratinizing, NOS |
| | Floor of mouth, NOS | Squamous cell carcinoma, NOS |
| | Gum, NOS | Squamous cell carcinoma, keratinizing, NOS |
| | Gum, NOS | Squamous cell carcinoma, NOS |
| | Hard palate | Squamous cell carcinoma, NOS |
| | Head, face or neck, NOS | Extra-adrenal paraganglioma, malignant |
| | Head, face or neck, NOS | Squamous cell carcinoma, NOS |
| | Hypopharynx, NOS | Squamous cell carcinoma, large cell, nonkeratinizing, NOS |
| | Hypopharynx, NOS | Squamous cell carcinoma, NOS |
| | Larynx, NOS | Basaloid squamous cell carcinoma |
| | Larynx, NOS | Squamous cell carcinoma, keratinizing, NOS |
| | Larynx, NOS | Squamous cell carcinoma, large cell, nonkeratinizing, NOS |
| | Larynx, NOS | Squamous cell carcinoma, NOS |
| | Lip, NOS | Squamous cell carcinoma, keratinizing, NOS |
| | Lip, NOS | Squamous cell carcinoma, NOS |
| | Lower gum | Squamous cell carcinoma, NOS |
| | Mandible | Squamous cell carcinoma, NOS |
| | Mouth, NOS | Squamous cell carcinoma, keratinizing, NOS |
| | Mouth, NOS | Squamous cell carcinoma, NOS |
| | Oropharynx, NOS | Squamous cell carcinoma, keratinizing, NOS |
| | Oropharynx, NOS | Squamous cell carcinoma, NOS |
| | Oropharynx, NOS | Squamous cell carcinoma, spindle cell |
| | Overlapping lesion of lip, oral cavity and pharynx | Squamous cell carcinoma, NOS |
| | Palate, NOS | Squamous cell carcinoma, keratinizing, NOS |
| | Pharynx, NOS | Squamous cell carcinoma, NOS |
| | Posterior wall of oropharynx | Squamous cell carcinoma, keratinizing, NOS |
| | Supraglottis | Squamous cell carcinoma, NOS |
| | Tongue, NOS | Leiomyosarcoma, NOS |
| | Tongue, NOS | Squamous cell carcinoma, keratinizing, NOS |
| | Tongue, NOS | Squamous cell carcinoma, NOS |
| | Tonsil, NOS | Basaloid squamous cell carcinoma |
| | Tonsil, NOS | Squamous cell carcinoma, keratinizing, NOS |
| | Tonsil, NOS | Squamous cell carcinoma, large cell, nonkeratinizing, NOS |
| | Tonsil, NOS | Squamous cell carcinoma, NOS |
| | Upper Gum | Squamous cell carcinoma, NOS |
| Hematological | Bone marrow | Acute myeloid leukemia, NOS |

| Cancer group | Histotype | Primary diagnosis |
|---|---|---|
| | Bone marrow | B lymphoblastic leukemia/lymphoma, NOS |
| | Bone marrow | Leukemia, NOS |
| | Bone marrow | Mixed phenotype acute leukemia with t(v;11q23); MLL rearranged |
| | Bone marrow | Mixed phenotype acute leukemia, B/myeloid, NOS |
| | Bone marrow | Mixed phenotype acute leukemia, T/myeloid, NOS |
| | Bone marrow | Multiple myeloma |
| | Bone marrow | Not Reported |
| | Bone marrow | Undifferentiated leukaemia |
| | Bones of skull and face and associated joints | Malignant lymphoma, large B-cell, diffuse, NOS |
| | Brain stem | Malignant lymphoma, large B-cell, diffuse, NOS |
| | Breast, NOS | Malignant lymphoma, large B-cell, diffuse, NOS |
| | Cerebellum, NOS | Malignant lymphoma, large B-cell, diffuse, NOS |
| | Connective, subcutaneous and other soft tissues of head, face, and neck | Malignant lymphoma, large B-cell, diffuse, NOS |
| | Connective, subcutaneous and other soft tissues, NOS | Malignant lymphoma, large B-cell, diffuse, NOS |
| | Intra-abdominal lymph nodes | Malignant lymphoma, large B-cell, diffuse, NOS |
| | Intrathoracic lymph nodes | Malignant lymphoma, large B-cell, diffuse, NOS |
| | Lymph nodes of axilla or arm | Malignant lymphoma, large B-cell, diffuse, NOS |
| | Lymph nodes of head, face and neck | Malignant lymphoma, large B-cell, diffuse, NOS |
| | Lymph nodes of inguinal region or leg | Malignant lymphoma, large B-cell, diffuse, NOS |
| | Specified parts of peritoneum | Malignant lymphoma, large B-cell, diffuse, NOS |
| | Submandibular gland | Malignant lymphoma, large B-cell, diffuse, NOS |
| | Thyroid gland | Malignant lymphoma, large B-cell, diffuse, NOS |
| Kidney | Kidney, NOS | Clear cell adenocarcinoma, NOS |
| | Kidney, NOS | Neuroblastoma, NOS |
| | Kidney, NOS | Papillary adenocarcinoma, NOS |
| | Kidney, NOS | Renal cell carcinoma, chromophobe type |
| | Kidney, NOS | Renal cell carcinoma, NOS |
| | Kidney, NOS | Synovial sarcoma, spindle cell |
| | Kidney, NOS | Wilms tumor |
| Liver and biliary tract | Ampulla of Vater | Adenocarcinoma, NOS |
| | Extrahepatic bile duct | Adenocarcinoma, NOS |
| | Extrahepatic bile duct | Cholangiocarcinoma |
| | Gallbladder | Cholangiocarcinoma |
| | Intrahepatic bile duct | Cholangiocarcinoma |
| | Liver | Cholangiocarcinoma |
| | Liver | Clear cell adenocarcinoma, NOS |

| Cancer group | Histotype | Primary diagnosis |
|---|---|---|
| | Liver | Combined hepatocellular carcinoma and cholangiocarcinoma |
| | Liver | Hepatocellular carcinoma, clear cell type |
| | Liver | Hepatocellular carcinoma, fibrolamellar |
| | Liver | Hepatocellular carcinoma, NOS |
| | Liver | Hepatocellular carcinoma, spindle cell variant |
| | Lower lobe, lung | Acinar cell carcinoma |
| | Lower lobe, lung | Adenocarcinoma with mixed subtypes |
| | Lower lobe, lung | Adenocarcinoma, NOS |
| | Lower lobe, lung | Basaloid squamous cell carcinoma |
| | Lower lobe, lung | Bronchio-alveolar carcinoma, mucinous |
| | Lower lobe, lung | Bronchiolo-alveolar adenocarcinoma, NOS |
| | Lower lobe, lung | Bronchiolo-alveolar carcinoma, non-mucinous |
| | Lower lobe, lung | Clear cell adenocarcinoma, NOS |
| | Lower lobe, lung | Micropapillary carcinoma, NOS |
| | Lower lobe, lung | Mucinous adenocarcinoma |
| | Lower lobe, lung | Papillary adenocarcinoma, NOS |
| | Lower lobe, lung | Papillary squamous cell carcinoma |
| | Lower lobe, lung | Solid carcinoma, NOS |
| | Lower lobe, lung | Squamous cell carcinoma, keratinizing, NOS |
| | Lower lobe, lung | Squamous cell carcinoma, large cell, nonkeratinizing, NOS |
| | Lower lobe, lung | Squamous cell carcinoma, NOS |
| | Lung, NOS | Adenocarcinoma with mixed subtypes |
| Lung and pleura | Lung, NOS | Adenocarcinoma, NOS |
| | Lung, NOS | Papillary adenocarcinoma, NOS |
| | Lung, NOS | Squamous cell carcinoma, keratinizing, NOS |
| | Lung, NOS | Squamous cell carcinoma, NOS |
| | Main bronchus | Adenocarcinoma, NOS |
| | Main bronchus | Squamous cell carcinoma, NOS |
| | Middle lobe, lung | Adenocarcinoma with mixed subtypes |
| | Middle lobe, lung | Adenocarcinoma, NOS |
| | Middle lobe, lung | Papillary adenocarcinoma, NOS |
| | Middle lobe, lung | Squamous cell carcinoma, keratinizing, NOS |
| | Middle lobe, lung | Squamous cell carcinoma, NOS |
| | Middle lobe, lung | Squamous cell carcinoma, small cell, nonkeratinizing |
| | Overlapping lesion of lung | Adenocarcinoma with mixed subtypes |
| | Overlapping lesion of lung | Bronchiolo-alveolar carcinoma, non-mucinous |
| | Overlapping lesion of lung | Papillary adenocarcinoma, NOS |
| | Overlapping lesion of lung | Squamous cell carcinoma, NOS |
| | Pleura, NOS | Epithelioid mesothelioma, malignant |
| | Pleura, NOS | Mesothelioma, biphasic, malignant |

| Cancer group | Histotype | Primary diagnosis |
|---|---|---|
| | Pleura, NOS | Mesothelioma, malignant |
| | Upper lobe, lung | Acinar cell carcinoma |
| | Upper lobe, lung | Adenocarcinoma with mixed subtypes |
| | Upper lobe, lung | Adenocarcinoma, NOS |
| | Upper lobe, lung | Basaloid squamous cell carcinoma |
| | Upper lobe, lung | Bronchio-alveolar carcinoma, mucinous |
| | Upper lobe, lung | Bronchiolo-alveolar adenocarcinoma, NOS |
| | Upper lobe, lung | Bronchiolo-alveolar carcinoma, non-mucinous |
| | Upper lobe, lung | Clear cell adenocarcinoma, NOS |
| | Upper lobe, lung | Micropapillary carcinoma, NOS |
| | Upper lobe, lung | Mucinous adenocarcinoma |
| | Upper lobe, lung | Papillary adenocarcinoma, NOS |
| | Upper lobe, lung | Papillary squamous cell carcinoma |
| | Upper lobe, lung | Signet ring cell carcinoma |
| | Upper lobe, lung | Solid carcinoma, NOS |
| | Upper lobe, lung | Squamous cell carcinoma, keratinizing, NOS |
| | Upper lobe, lung | Squamous cell carcinoma, large cell, nonkeratinizing, NOS |
| | Upper lobe, lung | Squamous cell carcinoma, NOS |
| Mediastinum | Anterior mediastinum | Malignant lymphoma, large B-cell, diffuse, NOS |
| | Anterior mediastinum | Thymic carcinoma, NOS |
| | Anterior mediastinum | Thymoma, type A, malignant |
| | Anterior mediastinum | Thymoma, type AB, malignant |
| | Anterior mediastinum | Thymoma, type AB, NOS |
| | Anterior mediastinum | Thymoma, type B1, malignant |
| | Anterior mediastinum | Thymoma, type B2, malignant |
| | Anterior mediastinum | Thymoma, type B2, NOS |
| | Anterior mediastinum | Thymoma, type B3, malignant |
| | Mediastinum, NOS | Extra-adrenal paraganglioma, malignant |
| | Mediastinum, NOS | Neuroblastoma, NOS |
| | Mediastinum, NOS | Thymoma, type A, malignant |
| | Mediastinum, NOS | Thymoma, type B1, malignant |
| | Mediastinum, NOS | Thymoma, type B2, malignant |
| | Posterior mediastinum | Neuroblastoma, NOS |
| | Thymus | Thymic carcinoma, NOS |
| | Thymus | Thymoma, type A, malignant |
| | Thymus | Thymoma, type A, NOS |
| | Thymus | Thymoma, type AB, malignant |
| | Thymus | Thymoma, type AB, NOS |
| | Thymus | Thymoma, type B1, malignant |
| | Thymus | Thymoma, type B1, NOS |
| | Thymus | Thymoma, type B2, malignant |
| | Thymus | Thymoma, type B2, NOS |

| Cancer group | Histotype | Primary diagnosis |
|---|---|---|
| | Thymus | Thymoma, type B3, malignant |
| Neuroendocrine | Abdomen, NOS | Ganglioneuroblastoma |
| | Abdomen, NOS | Neuroblastoma, NOS |
| | Aortic body and other paraganglia | Pheochromocytoma, malignant |
| | Bone marrow | Neuroblastoma, NOS |
| | Bones of skull and face and associated joints | Neuroblastoma, NOS |
| | Connective, subcutaneous and other soft tissues of abdomen | Ganglioneuroblastoma |
| | Connective, subcutaneous and other soft tissues of abdomen | Neuroblastoma, NOS |
| | Connective, subcutaneous and other soft tissues of abdomen | Paraganglioma, NOS |
| | Head, face or neck, NOS | Neuroblastoma, NOS |
| | Heart | Extra-adrenal paraganglioma, NOS |
| | Intra-abdominal lymph nodes | Neuroblastoma, NOS |
| | Liver | Neuroblastoma, NOS |
| | Long bones of lower limb and associated joints | Neuroblastoma, NOS |
| | Nervous system, NOS | Paraganglioma, malignant |
| | Pelvic lymph nodes | Ganglioneuroblastoma |
| | Peripheral nerves and autonomic nervous system of abdomen | Neuroblastoma, NOS |
| | Peripheral nerves and autonomic nervous system of thorax | Neuroblastoma, NOS |
| | Spinal meninges | Neuroblastoma, NOS |
| | Thorax, NOS | Extra-adrenal paraganglioma, NOS |
| | Thorax, NOS | Neuroblastoma, NOS |
| | Unknown | Ganglioneuroblastoma |
| | Unknown | Neuroblastoma, NOS |
| Ovary | Ovary | Cystadenocarcinoma, NOS |
| | Ovary | Leiomyosarcoma, NOS |
| | Ovary | Papillary serous cystadenocarcinoma |
| | Ovary | Serous cystadenocarcinoma, NOS |
| Pancreas | Body of pancreas | Adenocarcinoma, NOS |
| | Body of pancreas | Infiltrating duct carcinoma, NOS |
| | Body of pancreas | Neuroendocrine carcinoma, NOS |
| | Head of pancreas | Adenocarcinoma with mixed subtypes |
| | Head of pancreas | Adenocarcinoma, NOS |
| | Head of pancreas | Infiltrating duct carcinoma, NOS |
| | Head of pancreas | Mucinous adenocarcinoma |
| | Head of pancreas | Neuroendocrine carcinoma, NOS |
| | Overlapping lesion of pancreas | Adenocarcinoma, NOS |
| | Overlapping lesion of pancreas | Infiltrating duct carcinoma, NOS |
| | Pancreas, NOS | Adenocarcinoma, NOS |
| | Pancreas, NOS | Infiltrating duct carcinoma, NOS |
| | Pancreas, NOS | Neuroendocrine carcinoma, NOS |
| | Tail of pancreas | Adenocarcinoma, NOS |

| Cancer group | Histotype | Primary diagnosis |
|---|---|---|
| | Tail of pancreas | Carcinoma, undifferentiated, NOS |
| | Tail of pancreas | Infiltrating duct carcinoma, NOS |
| | Tail of pancreas | Neuroendocrine carcinoma, NOS |
| | Unknown primary site | Mucinous adenocarcinoma |
| Prostate | Prostate gland | Adenocarcinoma with mixed subtypes |
| | Prostate gland | Adenocarcinoma, NOS |
| | Prostate gland | Infiltrating duct carcinoma, NOS |
| | Prostate gland | Mucinous adenocarcinoma |
| Retroperitoneum | Overlapping lesion of retroperitoneum and peritoneum | Neuroblastoma, NOS |
| | Retroperitoneum | Abdominal fibromatosis |
| | Retroperitoneum | Dedifferentiated liposarcoma |
| | Retroperitoneum | Extra-adrenal paraganglioma, malignant |
| | Retroperitoneum | Extra-adrenal paraganglioma, NOS |
| | Retroperitoneum | Fibromyxosarcoma |
| | Retroperitoneum | Ganglioneuroblastoma |
| | Retroperitoneum | Giant cell sarcoma |
| | Retroperitoneum | Leiomyosarcoma, NOS |
| | Retroperitoneum | Liposarcoma, well differentiated |
| | Retroperitoneum | Myxoid leiomyosarcoma |
| | Retroperitoneum | Neuroblastoma, NOS |
| | Retroperitoneum | Undifferentiated sarcoma |
| | Specified parts of peritoneum | Dedifferentiated liposarcoma |
| Skin | Choroid | Epithelioid cell melanoma |
| | Choroid | Malignant melanoma, NOS |
| | Choroid | Mixed epithelioid and spindle cell melanoma |
| | Choroid | Spindle cell melanoma, NOS |
| | Choroid | Spindle cell melanoma, type B |
| | Ciliary body | Epithelioid cell melanoma |
| | Ciliary body | Mixed epithelioid and spindle cell melanoma |
| | Ciliary body | Spindle cell melanoma, NOS |
| | Overlapping lesion of eye and adnexa | Mixed epithelioid and spindle cell melanoma |
| | Overlapping lesion of eye and adnexa | Spindle cell melanoma, NOS |
| | Overlapping lesion of eye and adnexa | Spindle cell melanoma, type B |
| | Skin, NOS | Acral lentiginous melanoma, malignant |
| | Skin, NOS | Amelanotic melanoma |
| | Skin, NOS | Desmoplastic melanoma, malignant |
| | Skin, NOS | Epithelioid cell melanoma |
| | Skin, NOS | Lentigo maligna melanoma |
| | Skin, NOS | Malignant melanoma, NOS |
| | Skin, NOS | Mixed epithelioid and spindle cell melanoma |
| | Skin, NOS | Nodular melanoma |

| Cancer group | Histotype | Primary diagnosis |
|---|---|---|
| | Skin, NOS | Spindle cell melanoma, NOS |
| | Skin, NOS | Spindle cell sarcoma |
| | Skin, NOS | Superficial spreading melanoma |
| Small & large bowel | Ascending colon | Adenocarcinoma, NOS |
| | Ascending colon | Adenosquamous carcinoma |
| | Ascending colon | Mucinous adenocarcinoma |
| | Ascending colon | Papillary adenocarcinoma, NOS |
| | Cecum | Adenocarcinoma, NOS |
| | Cecum | Mucinous adenocarcinoma |
| | Colon, NOS | Adenocarcinoma with neuroendocrine differentiation |
| | Colon, NOS | Adenocarcinoma, NOS |
| | Colon, NOS | Malignant lymphoma, large B-cell, diffuse, NOS |
| | Colon, NOS | Mucinous adenocarcinoma |
| | Descending colon | Adenocarcinoma, NOS |
| | Descending colon | Dedifferentiated liposarcoma |
| | Descending colon | Mucinous adenocarcinoma |
| | Hepatic flexure of colon | Adenocarcinoma, NOS |
| | Hepatic flexure of colon | Mucinous adenocarcinoma |
| | Jejunum | Malignant lymphoma, large B-cell, diffuse, NOS |
| | Rectosigmoid junction | Adenocarcinoma, NOS |
| | Rectosigmoid junction | Mucinous adenocarcinoma |
| | Rectum, NOS | Adenocarcinoma in tubolovillous adenoma |
| | Rectum, NOS | Adenocarcinoma with mixed subtypes |
| | Rectum, NOS | Adenocarcinoma, NOS |
| | Rectum, NOS | Mucinous adenocarcinoma |
| | Rectum, NOS | Tubular adenocarcinoma |
| | Sigmoid colon | Adenocarcinoma, NOS |
| | Sigmoid colon | Carcinoma, NOS |
| | Sigmoid colon | Mucinous adenocarcinoma |
| | Sigmoid colon | Papillary adenocarcinoma, NOS |
| | Small intestine, NOS | Malignant lymphoma, large B-cell, diffuse, NOS |
| | Splenic flexure of colon | Adenocarcinoma, NOS |
| | Splenic flexure of colon | Mucinous adenocarcinoma |
| | Transverse colon | Adenocarcinoma, NOS |
| | Transverse colon | Mucinous adenocarcinoma |
| Spermatic cord | Spermatic cord | Dedifferentiated liposarcoma |
| Stomach | Body of stomach | Adenocarcinoma, intestinal type |
| | Body of stomach | Adenocarcinoma, NOS |
| | Body of stomach | Carcinoma, diffuse type |
| | Body of stomach | Mucinous adenocarcinoma |
| | Body of stomach | Papillary adenocarcinoma, NOS |
| | Body of stomach | Signet ring cell carcinoma |

| Cancer group | Histotype | Primary diagnosis |
|---|---|---|
| | Body of stomach | Tubular adenocarcinoma |
| | Cardia, NOS | Adenocarcinoma, intestinal type |
| | Cardia, NOS | Adenocarcinoma, NOS |
| | Cardia, NOS | Carcinoma, diffuse type |
| | Cardia, NOS | Mucinous adenocarcinoma |
| | Cardia, NOS | Papillary adenocarcinoma, NOS |
| | Cardia, NOS | Signet ring cell carcinoma |
| | Cardia, NOS | Squamous cell carcinoma, NOS |
| | Cardia, NOS | Tubular adenocarcinoma |
| | Fundus of stomach | Adenocarcinoma, intestinal type |
| | Fundus of stomach | Adenocarcinoma, NOS |
| | Fundus of stomach | Carcinoma, diffuse type |
| | Fundus of stomach | Mucinous adenocarcinoma |
| | Fundus of stomach | Papillary adenocarcinoma, NOS |
| | Fundus of stomach | Tubular adenocarcinoma |
| | Gastric antrum | Adenocarcinoma, intestinal type |
| | Gastric antrum | Adenocarcinoma, NOS |
| | Gastric antrum | Carcinoma, diffuse type |
| | Gastric antrum | Mucinous adenocarcinoma |
| | Gastric antrum | Papillary adenocarcinoma, NOS |
| | Gastric antrum | Signet ring cell carcinoma |
| | Gastric antrum | Tubular adenocarcinoma |
| | Lesser curvature of stomach, NOS | Adenocarcinoma, NOS |
| | Stomach, NOS | Adenocarcinoma with mixed subtypes |
| | Stomach, NOS | Adenocarcinoma, intestinal type |
| | Stomach, NOS | Adenocarcinoma, NOS |
| | Stomach, NOS | Carcinoma, diffuse type |
| | Stomach, NOS | Dedifferentiated liposarcoma |
| | Stomach, NOS | Leiomyosarcoma, NOS |
| | Stomach, NOS | Malignant lymphoma, large B-cell, diffuse, NOS |
| | Stomach, NOS | Mucinous adenocarcinoma |
| Testis | Testis, NOS | Embryonal carcinoma, NOS |
| | Testis, NOS | Malignant lymphoma, large B-cell, diffuse, NOS |
| | Testis, NOS | Mixed germ cell tumor |
| | Testis, NOS | Seminoma, NOS |
| | Testis, NOS | Teratocarcinoma |
| | Testis, NOS | Teratoma, benign |
| | Testis, NOS | Teratoma, malignant, NOS |
| | Testis, NOS | Yolk sac tumor |
| Thyroid | Thyroid gland | Carcinoma, NOS |
| | Thyroid gland | Follicular adenocarcinoma, NOS |
| | Thyroid gland | Follicular carcinoma, minimally invasive |
| | Thyroid gland | Nonencapsulated sclerosing carcinoma |

| Cancer group | Histotype | Primary diagnosis |
|---|---|---|
| | Thyroid gland | Oxyphilic adenocarcinoma |
| | Thyroid gland | Papillary adenocarcinoma, NOS |
| | Thyroid gland | Papillary carcinoma, columnar cell |
| | Thyroid gland | Papillary carcinoma, follicular variant |
| Urinary tract | Anterior wall of bladder | Papillary transitional cell carcinoma |
| | Anterior wall of bladder | Transitional cell carcinoma |
| | Bladder neck | Transitional cell carcinoma |
| | Bladder, NOS | Carcinoma, NOS |
| | Bladder, NOS | Papillary transitional cell carcinoma |
| | Bladder, NOS | Squamous cell carcinoma, NOS |
| | Bladder, NOS | Transitional cell carcinoma |
| | Dome of bladder | Papillary adenocarcinoma, NOS |
| | Dome of bladder | Transitional cell carcinoma |
| | Lateral wall of bladder | Papillary transitional cell carcinoma |
| | Lateral wall of bladder | Transitional cell carcinoma |
| | Posterior wall of bladder | Papillary transitional cell carcinoma |
| | Posterior wall of bladder | Transitional cell carcinoma |
| | Trigone of bladder | Papillary transitional cell carcinoma |
| | Trigone of bladder | Transitional cell carcinoma |
| | Ureteric orifice | Transitional cell carcinoma |
| Uterus | Cervix uteri | Adenocarcinoma, endocervical type |
| | Cervix uteri | Adenocarcinoma, NOS |
| | Cervix uteri | Adenosquamous carcinoma |
| | Cervix uteri | Basaloid squamous cell carcinoma |
| | Cervix uteri | Endometrioid adenocarcinoma, NOS |
| | Cervix uteri | Mucinous adenocarcinoma, endocervical type |
| | Cervix uteri | Papillary squamous cell carcinoma |
| | Cervix uteri | Squamous cell carcinoma, keratinizing, NOS |
| | Cervix uteri | Squamous cell carcinoma, large cell, nonkeratinizing, NOS |
| | Cervix uteri | Squamous cell carcinoma, NOS |
| | Corpus uteri | Carcinosarcoma, NOS |
| | Corpus uteri | Endometrioid adenocarcinoma, NOS |
| | Corpus uteri | Leiomyosarcoma, NOS |
| | Corpus uteri | Mullerian mixed tumor |
| | Endometrium | Adenocarcinoma, NOS |
| | Endometrium | Carcinoma, undifferentiated, NOS |
| | Endometrium | Clear cell adenocarcinoma, NOS |
| | Endometrium | Endometrioid adenocarcinoma, NOS |
| | Endometrium | Endometrioid adenocarcinoma, secretory variant |
| | Endometrium | Papillary serous cystadenocarcinoma |
| | Endometrium | Serous cystadenocarcinoma, NOS |
| | Endometrium | Serous surface papillary carcinoma |

| Cancer group | Histotype | Primary diagnosis |
|---|---|---|
| | Fundus uteri | Endometrioid adenocarcinoma, NOS |
| | Fundus uteri | Serous cystadenocarcinoma, NOS |
| | Isthmus uteri | Endometrioid adenocarcinoma, NOS |
| | Isthmus uteri | Serous cystadenocarcinoma, NOS |
| | Myometrium | Leiomyosarcoma, NOS |
| | Uterus, NOS | Carcinosarcoma, NOS |
| | Uterus, NOS | Leiomyosarcoma, NOS |
| | Uterus, NOS | Mesodermal mixed tumor |
| | Uterus, NOS | Mullerian mixed tumor |
| | Uterus, NOS | Myxoid leiomyosarcoma |

**Supplementary Table 2: Median survival and hazard ratios for the Kaplan Meier curves shown in Figure 2.**

Statistical significance was evaluated using the log-rank (Mantel-Cox) test. n, number of patients in the cluster; MOS, median OS (in years); CI95, 95% confidence interval; HR, hazard ratio; NR, not reachable.

| Cancer group | Cluster 0 | | | Cluster 1 | | | Comparative | | |
|---|---|---|---|---|---|---|---|---|---|
| | n | MOS | MS CI95 | n | MOS | MS CI95 | p-value | HR | HR CI95 |
| Small & large bowel | 30 | 3.2 | 1.49-NR | 523 | 8.3 | 5.84-NR | .004 | 2.5 | 1.33-4.76 |
| Lung & pleura | 149 | 2.9 | 2.42-3.72 | 1079 | 4.4 | 3.69-4.87 | .003 | 1.47 | 1.14-1.89 |
| Urinary tract | 30 | 1.6 | 0.97-NR | 370 | 2.9 | 2.13-5.4 | .045 | 1.67 | 1.01-2.78 |
| Adrenal | 218 | NR | NR | 111 | 4.8 | 3.85-7.79 | .0002 | 2.13 | 1.43-3.23 |
| CNS | 187 | 4.1 | 2.8-12.08 | 784 | 2.0 | 1.77-2.2 | .00003 | 1.75 | 1.33-2.27 |
| Mediastinum | 89 | NR | NR | 37 | NR | 7.97-NR | .036 | 3.45 | 1.09-11.11 |

**Supplementary Table 3: Pathologies included in the 24 selected cancer groups.**

Pathologies are defined according to either the tumor type or to the precise anatomic location. This definition only includes those pathologies for which a relevant number of patients were found in the TCGA dataset, and it is intended to be as inclusive as possible, hence overlapping criteria have been followed in some cases. Therefore, some patients may have been included in more than one pathology.

(1) These pathologies are paraganglioma, ganglioneuroblastoma and neuroblastoma

(2) MPNST, malignant peripheral nerve sheath tumor

(3) aNSCLC, adenocarcinoma non-small cell lung cancer

(4) eNSCLC, epidermoid non-small cell lung cancer

| Cancer group | Pathologies |
|---|---|
| Adrenal | Adrenal carcinoma |
| | Pheochromocytoma |
| | Neuroblastoma |
| | Adrenal cancer of neural origin (1) |
| | Adrenocortical carcinoma |
| | Adrenal medullary carcinoma |
| Bone | Osteosarcoma |
| Breast | Ductal breast carcinoma |
| | Lobular breast carcinoma |
| | Breast cancer - other |
| CNS | Anaplasic astrocytoma |
| | Non-anaplasic astrocytoma |
| | Glioblastoma |
| | Oligodendroglioma |
| | CNS - other |
| Connective & soft tissue | Sarcoma |
| | Histiocytoma |
| | MPNST (2) |
| Esophagus | Esophagous adenocarcinoma |
| | Esophagus squamous cell carcinoma |
| | Middle esophageal carcinoma |
| | Lower esophageal carcinoma |
| Head & neck | Mouth cancer |
| | Pharynx cancer |
| | Larynx cancer |
| | Tongue cancer |
| Hematological | Acute lymphocitic leukemia |
| | Acute myeloid leukemia |
| | Lymphoma |
| | Myeloma |
| Kidney | Clear cell renal cell carcinoma |
| | Chromophobe renal cell carcinoma |

| Cancer group | Pathologies |
| --- | --- |
| | Papilary renal cell carcinoma |
| | Nephroblastoma |
| | Kidney cancer - other |
| Liver and biliary tract | Liver adenocarcinoma |
| | Hepatocellular carcinoma |
| | Cholangiocarcinoma |
| Lung and pleura | Mesothelioma |
| | Lung adenocarcinoma (aNSCLC)[(3)] |
| | Squamous cell lung cancer (eNSCLC)[(4)] |
| Mediastinum | Mediastinal epithelial neoplasia |
| | Anterior mediastinum tumor |
| | Thymoma |
| Neuroendocrine | Neuroblastoma |
| | Other - neuroendocrine |
| Ovary | Ovarian cancer |
| Pancreas | Exocrine pancreatic cancer |
| | Pancreatic adenocarcinoma |
| | Pancreatic ductal adenocarcinoma |
| Retroperitoneum | Retroperitoneal sarcoma |
| Prostate | Prostate adenocarcinoma |
| Skin | Skin melanoma |
| | Ocular melanoma |
| Small & large bowel | Colorectal cancer |
| | Rectal cancer |
| | Colon cancer |
| | Right colon cancer |
| | Left colon cancer |
| | Colorectal adenocarcinoma |
| | Mucinous colorectal adenocarcinoma |
| Stomach | Gastric adenocarcinoma |
| | Gastric diffuse adenocarcinoma |
| | Intestinal type gastric adenocarcinoma |
| Testis | Testicular cancer |
| | Testicular germ cell cancer |
| Thyroid | Thyroid cancer |
| Urinary tract | Urothelial carcinoma |
| Uterus | Cervical cancer |
| | Endometrial cancer |
| | Cancer of the corpus uteri |
| | Uterine serous carcinoma |
| | Uterine squamous cell carcinoma |
| | Uterine adenocarcinoma |
| | Endometrial adenocarcinoma |
| | Mixed Mullerian tumor |

**Supplementary Table 4: List of genes harboring mutations exclusively found in non-metastatic CRC patients included in cluster 0**

| Genes name |
|---|
| OR5C1 |
| LOC107985532 |
| MAP7D2 |
| SNORD116-29 |
| NMS |
| GOLGA7 |
| SNORD3B-1 |
| LINC01098 |
| NKX2-4 |
| MIR17HG |
| IGF2-AS |
| GIP |
| CBWD6 |
| MIR889 |
| CCDC163 |

**Supplementary Table 5: List of genes harboring mutations exclusively found in stage-1 eNSCLC patients included in cluster 0**

| Genes names | | | | |
|---|---|---|---|---|
| ABT1 | CTNS | HMGB3 | PANK2 | ST6GALNAC4 |
| ACTL6A | CUZD1 | HOPX | PEA15 | STH |
| AHCYL1 | CXCL13 | HOXB6 | PHB2 | STMN4 |
| ALOX12 | DAZAP1 | HSD17B3 | PHF2 | SUCLG1 |
| ANKRD46 | DCSTAMP | IGF2-AS | PLA2G2A | SUPT7L |
| APOC1 | DCUN1D4 | KLF11 | PRAMEF5 | SYT15 |
| ARNTL | DHX29 | KRTAP6-2 | PROSER2 | TAF5 |
| ASAP3 | DIP2C-AS1 | LAPTM4A | PRR22 | TALDO1 |
| ATP6V1E2 | DNAJC17 | LCE2A | PRSS54 | TIGD6 |
| AZGP1 | DPH5 | LIPN | PTX4 | TM2D3 |
| BCL2L1 | EEF1A1 | LRFN3 | RAB37 | TMCO2 |
| BCL2L2 | EEF1AKMT4 | LRRC75A | RAB4A | TMEM256 |
| BIVM | EFNA1 | LSM11 | RBM34 | TMEM99 |
| C16orf82 | EFNB1 | LYSMD4 | RELB | TMX1 |
| C19orf48 | EIF4E3 | MAP2 | RETSAT | TSPAN17 |
| C1orf50 | ENAH | MBNL1 | ROPN1B | TSTD2 |
| C1R | FAM90A10P | MED14 | RPL23 | TWSG1 |
| C22orf23 | FAM91A1 | MEOX1 | S100A11 | UBE2J2 |
| C9orf47 | FAM9B | METAP1 | SH3BGRL | UBE2M |
| C9orf72 | FAXDC2 | MIR103A2 | SKA3 | UGT1A1 |
| CA1 | FOSB | MIR1291 | SLC25A14 | UGT1A7 |
| CALM2 | FUT2 | MIR4713HG | SLC25A47 | UPK1A |
| CCK | FXYD5 | MIR585 | SLC46A3 | UQCR10 |
| CCL26 | GABARAP | MOK | SNAPC2 | VPS37A |
| CDC20 | GAR1 | MREG | SNORD115-15 | WNT9B |
| CDC42EP1 | GFRA3 | MROH8 | SNORD116-14 | XKR6 |
| CDK20 | GPR135 | MRPL10 | SNORD116-2 | YEATS4 |
| CES4A | GUK1 | MTCH2 | SNORD33 | ZBTB5 |
| CHCHD6 | H2BC9 | MZB1 | SNORD82 | ZNF18 |
| CHP2 | H3C8 | NELFA | SNURF | ZNF445 |
| CKAP2 | HAGHL | NFKBIA | SP3 | ZNF581 |
| CORO6 | HCAR1 | NHLRC1 | SSX5 | ZNF654 |
| CRBN | HLA-DPA1 | OR4F17 | ST3GAL6 | ZRSR2 |
| CRYBB2 | HMGB2 | | | |

**Supplementary Table 6: List of genes harboring mutations exclusively found in pheochromocytoma patients included in cluster 0**

| Genes names | | | | |
|---|---|---|---|---|
| ABCA12 | DHTKD1 | JOSD1 | OR51V1 | SLC4A2 |
| ABCA2 | DOCK7 | KAT6A | OR52I2 | SNX19 |
| ABCD1 | DPP9 | KBTBD2 | OSBPL6 | SOX5 |
| ABCD3 | DSC2 | KCTD3 | OTOP1 | SPDL1 |
| ADAM2 | DST | KDM6B | OTOP3 | SPEN |
| ADCY10 | DUOX2 | KIAA0319 | PADI3 | SPINK5 |
| ADGRG6 | DVL2 | KLF12 | PAIP1 | SRP72 |
| AK9 | DYSF | KLHDC3 | PAK3 | SSX6P |
| AKAP13 | ECHDC2 | KLHL36 | PAPOLA | ST6GAL2 |
| ALDH2 | EGLN1 | KLRD1 | PCK1 | STXBP3 |
| ALKBH1 | EHMT1 | KRTAP6-2 | PGR | SUPT16H |
| AOC3 | EIF4G3 | KSR1 | PHKA2 | SYT3 |
| AP4E1 | ENOX2 | LANCL1 | PHTF2 | SYT6 |
| APC | EPC1 | LARP4B | PIGO | TAB3 |
| APOL2 | EPHA3 | LATS2 | PKN1 | TACC2 |
| ARAP3 | EPOR | LCT | PNO1 | TAF1L |
| ARHGEF39 | ERCC6L2 | LINC00221 | POLR2A | TBC1D1 |
| ARHGEF40 | ESX1 | LLGL1 | POLR3B | TBC1D10C |
| ARID2 | F5 | LPAR4 | PPP1R10 | TBP |
| ARID3C | FAM122C | LSM11 | PPT2 | TCAP |
| ARL2 | FAM83D | LTBP1 | PRG2 | TDRD9 |
| ASB9 | FBXL20 | MAGEA12 | PRICKLE2 | TEC |
| ASRGL1 | FDXR | MAGEB1 | PRKACB | TFR2 |
| ATP2A1 | FGF16 | MALT1 | PRKAR1A | THBS1 |
| ATP8B2 | FLII | MAML3 | PRKD1 | THOC1 |
| ATP9A | FMO3 | MAN1B1 | PROM1 | THSD7B |
| ATR | FOXA2 | MARK4 | PRRC2C | TM2D1 |
| ATRX | FOXI2 | MATR3 | PSD4 | TM9SF4 |
| BCAS1 | FOXM1 | MBD5 | PTPN5 | TMEM208 |
| BCL2L2 | FZD8 | MCOLN2 | PVALB | TMEM8B |
| BCR | GATAD2B | MERTK | PYGB | TNKS |
| BICD1 | GBP6 | MIR4436A | QPCT | TOX3 |
| BPIFB6 | GJA5 | MIR937 | R3HDML | TPCN1 |
| BRWD3 | GLB1L3 | MLXIP | RAB35 | TPO |
| BTBD10 | GLOD4 | MMP2 | RABEPK | TRIM17 |
| BZW2 | GOLGA4 | MMRN1 | RAD50 | TRIP11 |

| Genes names | | | | |
|---|---|---|---|---|
| C5 | GOLM2 | MRC2 | REPIN1 | TRIP6 |
| CACYBP | GOSR2 | MRGPRX3 | ROCK2 | TRRAP |
| CALCRL | GP2 | MSLN | RP1L1 | TUB |
| CASP2 | GPR137 | MUC5AC | RPS27A | TXNRD1 |
| CCDC110 | GPR148 | MYH15 | RPS6KA5 | UBN1 |
| CCDC171 | GPR156 | MYO5C | RPS6KC1 | UNC5A |
| CCDC60 | GPX4 | MYPN | RRBP1 | URM1 |
| CCL13 | GTF2H2 | NAV3 | RSF1 | USP19 |
| CDH20 | GULP1 | NBN | RTL9 | VAV1 |
| CDH5 | H1-0 | NCAPD3 | RTP4 | VPS13C |
| CDK5RAP1 | HAPLN4 | NEK4 | RWDD4 | WDR62 |
| CDYL2 | HAT1 | NFIC | SALL1 | WDR64 |
| CELSR3 | HAVCR1 | NIBAN1 | SCN5A | WNT3 |
| CEP250 | HDHD5 | NLRP4 | SCRIB | WT1 |
| CFH | HECA | NOL4 | SEC14L3 | XPNPEP2 |
| CHST15 | HECTD1 | NOP58 | SEC14L5 | YME1L1 |
| CLASP2 | HELZ | NR1H4 | SH3BP4 | ZACN |
| COASY | HFM1 | NRROS | SHB | ZBTB14 |
| COL12A1 | HSP90AA1 | NSD2 | SHISA3 | ZNF275 |
| CPSF4 | IGLL1 | NUCKS1 | SHPRH | ZNF507 |
| CRAMP1 | IL15RA | NUGGC | SIMC1 | ZNF516 |
| CSH2 | IL34 | NUP98 | SLC12A4 | ZNF613 |
| CTH | IQSEC3 | NUTM1 | SLC12A8 | ZNF676 |
| CYP2B6 | IRF9 | OLIG3 | SLC26A9 | ZNF687 |
| CYP4F2 | ITLN2 | OPRPN | SLC35C1 | ZNF799 |
| DCAF11 | IVL | OR10T2 | SLC38A4 | ZZEF1 |
| DGAT2 | IWS1 | OR10X1 | | |

**Supplementary Table 7: List of 75 genes belonging to the PI3K/Akt signaling pathway that were selected from breast cancer patients included in SP1 after the triage described in the text.**

| **Genes names** | | | | |
|---|---|---|---|---|
| ANGPT4 | CRTC2 | IRS1 | LAMB4 | PPP2R5B |
| ATF2 | CSF1 | ITGA1 | LAMC1 | PPP2R5C |
| CCNE2 | EFNA3 | ITGA11 | LAMC2 | PPP2R5D |
| COL1A2 | EFNA5 | ITGA7 | LAMC3 | PRLR |
| COL4A1 | FGF18 | ITGA9 | LPAR1 | RELN |
| COL4A2 | FGF6 | ITGB4 | LPAR5 | RHEB |
| COL4A3 | FGF8 | ITGB5 | MAGI1 | RPS6 |
| COL4A4 | FN1 | ITGB6 | PCK1 | RPS6KB2 |
| COL4A6 | GNB3 | LAMA1 | PCK2 | SGK2 |
| COL6A2 | GNB5 | LAMA2 | PDGFC | SPP1 |
| COL6A3 | GYS1 | LAMA3 | PHLPP1 | THBS3 |
| COL9A1 | HGF | LAMA4 | PHLPP2 | TNN |
| COL9A3 | IFNA16 | LAMA5 | PPP2CB | TNXB |
| COMP | IFNA2 | LAMB1 | PPP2R2B | VEGFD |
| CREB3L3 | IFNA21 | LAMB2 | PPP2R3C | YWHAB |
